# Perisaccadic and Attentional Remapping of Receptive Fields in Lateral Intraparietal Area and Frontal Eye Fields

**DOI:** 10.1101/2023.09.23.558993

**Authors:** Xiao Wang, Cong Zhang, Lin Yang, Min Jin, Michael E. Goldberg, Mingsha Zhang, Ning Qian

**Affiliations:** State Key Laboratory of Cognitive Neuroscience and Learning IDG/McGovern Institute for Brain Research, Beijing Normal University Beijing, China; Department of Neuroscience and Zuckerman Institute, Columbia University, New York, NY, USA; Departments of Neurology, Psychiatry, and Ophthalmology, Columbia University, New York, NY, USA; Department of Physiology & Cellular Biophysics Columbia University, New York, NY, USA

**Keywords:** visuomotor interactions, efference copy, space constancy, working memory, coordinate transformation, compressive mislocalization

## Abstract

The nature and function of perisaccadic receptive-field (RF) remapping have been controversial. We used a delayed saccade task to reduce previous confounds and examined the remapping time course in areas LIP and FEF. In the delay period, the RF shift direction turned from the initial fixation to the saccade target. In the perisaccadic period, RFs first shifted toward the target (convergent remapping) but around the time of saccade onset/offset, the shifts became predominantly toward the post-saccadic RF locations (forward remapping). Thus, unlike forward remapping that depends on the corollary discharge (CD) of the saccade command, convergent remapping appeared to follow attention from the initial fixation to the target. We modelled the data with attention-modulated and CD-gated connections, and showed that both sets of connections emerged automatically in neural networks trained to update stimulus retinal locations across saccades. Our work thus unifies previous findings into a mechanism for transsaccadic visual stability.

## Introduction

Visual information enters the brain via the retina, which projects a point-to-point retinotopic map to the visual cortex via the lateral geniculate. The retinotopic map is maintained throughout a number of prestriate visual areas ^1,2^. However, a retinotopic map is inadequate for spatially accurate perception, because eye movements change the retinal location of a given object in the world. Nonetheless the spatial perception of humans and monkeys is largely independent of gaze. A classic demonstration of the brain’s ability to convert a retinotopic map into a spatially accurate map is the double-step task ^3,4^. Subjects must make successive saccades to two flashed targets both of which disappear before the first saccade. The retinal and oculomotor vectors of the first saccade are identical. However, the first saccade creates a dissonance between the spatial and retinotopic locations of the second saccade target, and the brain must compensate for this dissonance. Helmholtz ^5^ theorized that the brain used the motor signal for the first saccade to update the visual representation and create a spatially accurate visual signal for perception and action. Duhamel et al ^6^ showed that Helmholtz’s theory has a physiological correlate. When monkeys fixate, a given object in space occupies the RF of a neuron, the current RF (cRF). When monkeys make a saccade, the saccade will move the RF to a new spatial location, the future RF (fRF), even though the object’s spatial location has not changed. Neurons in the lateral intraparietal area (LIP) respond to a stimulus in the fRF even before the saccade, effecting a forward shift of the RF in the direction of the saccade. Because the shift starts before the eye moves, the signal causing the shift must arise from a motor signal feeding back to the sensory system, a phenomenon now known as corollary discharge (CD). This forward shift is found in many brain areas, including the superior colliculus (SC) ^7^, the frontal eye fields (FEF) ^8^, V3 ^9^, and the parietal reach area ^10^. In LIP the perisaccadic visual responses are not limited to the cRF and fRF. Instead, stimuli positioned in any of the spatial locations across which the saccade will sweep the retinal RF will evoke a response before the saccade (Fig. 1a). Stimuli closer to the fRF evoke larger responses with longer latencies than stimuli closer to the cRF ^11^. Forward remapping has been postulated to provide a key mechanism for the transsaccadic maintenance of a spatially accurate signal ^12^. Without forward remapping the brain’s representation of the visual world would be inaccurate for at least 45 ms ^13^, and behaviorally relevant visual signals inaccurate for 90 ms ^14^ after every saccade.

**Fig. 1.**
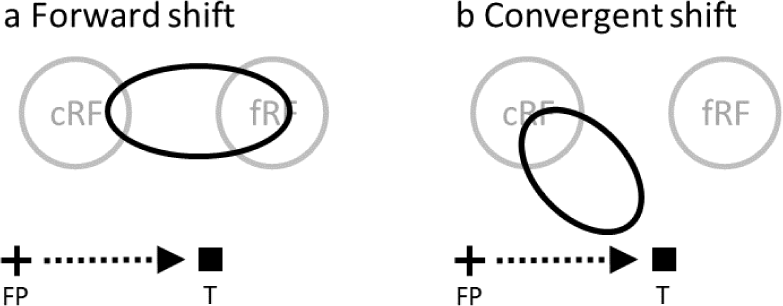
Perisaccadic RF remapping found in LIP, FEF, and other brain areas, drawn in a screen reference frame. The cross, square and arrow represent the fixation point (FP), saccade target (T), and saccade vector, respectively. cRF and fRF refer to a cell’s current (pre-saccadic) and future (post-saccadic) RFs, respectively. In each panel, the region(s) enclosed by black oval represent perisaccadic RF. (a) Forward shift in the saccade direction. (b) Convergent shift toward the target.

The original remapping studies only probed a cell’s responses at a few positions. To measure perisaccadic RFs more completely, an influential study ^15^ sampled FEF cells’ responses from an array of spatial positions. The study found that around saccade onset, RFs shift toward the saccade target (Fig. 1b), instead of toward the fRFs. The conclusion was that this convergent shift is a substrate of attention to the saccade target, unrelated to the maintenance of a spatially accurate signal across saccades, despite the neuropsychological evidence. The study argued that the forward shift may be an artifact of under-sampling the probe locations. However, the methods used caused a number of confounds: 1) It is well known that both the abrupt onset of visual stimuli ^16^ and the saccade target ^17^ evoke attention, and RFs shift toward the locus of attention ^18^. Most remapping studies, including that of Zirnsak et al, use the onset of the saccade target as the saccade go signal, and thus could not separate the effect of the attention evoked by the target onset from that of the saccade CD on the RF shifts. 2) The study integrated neuronal activities from 50 to 350 ms after the probe onset. This large time window must average attentional and CD effects together, and the strong attention from the target onset could mask the CD effect. Additionally, around the saccade onset, neurons in LIP (one synapse away from the FEF) exhibit progressive RF shifts from cRF to fRF ^11^, encompassing the entire portion of the space between them (Fig. 1a). The large time window could average response from the enlarged receptive field and underestimate the maximum forward-shift amplitude. 3) There might be differences between LIP and FEF.

Here we used a delayed saccade task to separate the attentional effect of the target onset and the effect of the saccade CD, and recorded LIP and FEF single-unit activities evoked by probes at different locations and times. The task allowed us to investigate the time course of the RF remapping in detail. We found that in the delay period, RFs shifted slightly in a direction close to the initial fixation 50 ms after the probe onset, but shifted toward the target 250 ms after the probe onset. In the perisaccadic period, RFs shifted toward the target 50 ms after the probe onset, but around the time of saccade onset/offset, the shifts became predominantly forward toward fRFs and the amplitudes approached that of the saccades. When we integrated neuronal activities from 50 to 350 ms after the probe onset, perisaccadic RFs were still closer to the fRFs than to the target, indicating stronger forward than convergent remapping. Since it first appeared in the delay period when the saccade was suppressed, convergent remapping is not really perisaccadic but attentional.

To explain our data, we constructed a circuit model by integrating CD-gated directional connections for forward remapping ^11^ and attention-modulated center/surround connections ^19,20^ for convergent remapping. We further demonstrated that both sets of connections emerged automatically in artificial neural networks trained to update retinal positions of stimuli across saccades. Mechanistically, the center/surround connections provide attractor dynamics to represent the retinal position of a stimulus as an activity bump, and the CD-gated connections move the activity bump for transsaccadic updating. The result has the surprising functional implication that the center/surround connections might not only be modulated by attention to produce convergent RF remapping but also work synergistically with the CD-gated connections to enable accurate spatial perception across saccades.

## Results

We used a delayed saccade task (Fig. 2) to sample a cell’s responses from a grid of spatial positions (tailored for each cell according to pilot RF mapping) and four time epochs (after the monkeys achieved initial fixation, after the appearance of the target, after the disappearance of the fixation (the go signal for the saccade), and well after the saccade, respectively). In a given trial we flashed one probe stimulus in each epoch at a random grid position, and across trials we sampled all grid positions and epochs. We name the four epochs pre-target (**c**urrent), **d**elay, **p**erisaccadic, and post-saccadic (**f**uture) periods (Fig. 2b), and denote a cell’s RFs mapped in these periods as its **c**RF, **d**RF, **p**RF, and **f**RF, respectively. We recorded a total of 391 and 427 single units from LIP and FEF, respectively, in 3 macaques. We then screened the data to select cells with significant visual responses, well-sampled RFs, and significant RF shifts in the delay or perisaccadic epoch (Methods).

**Fig. 2.**
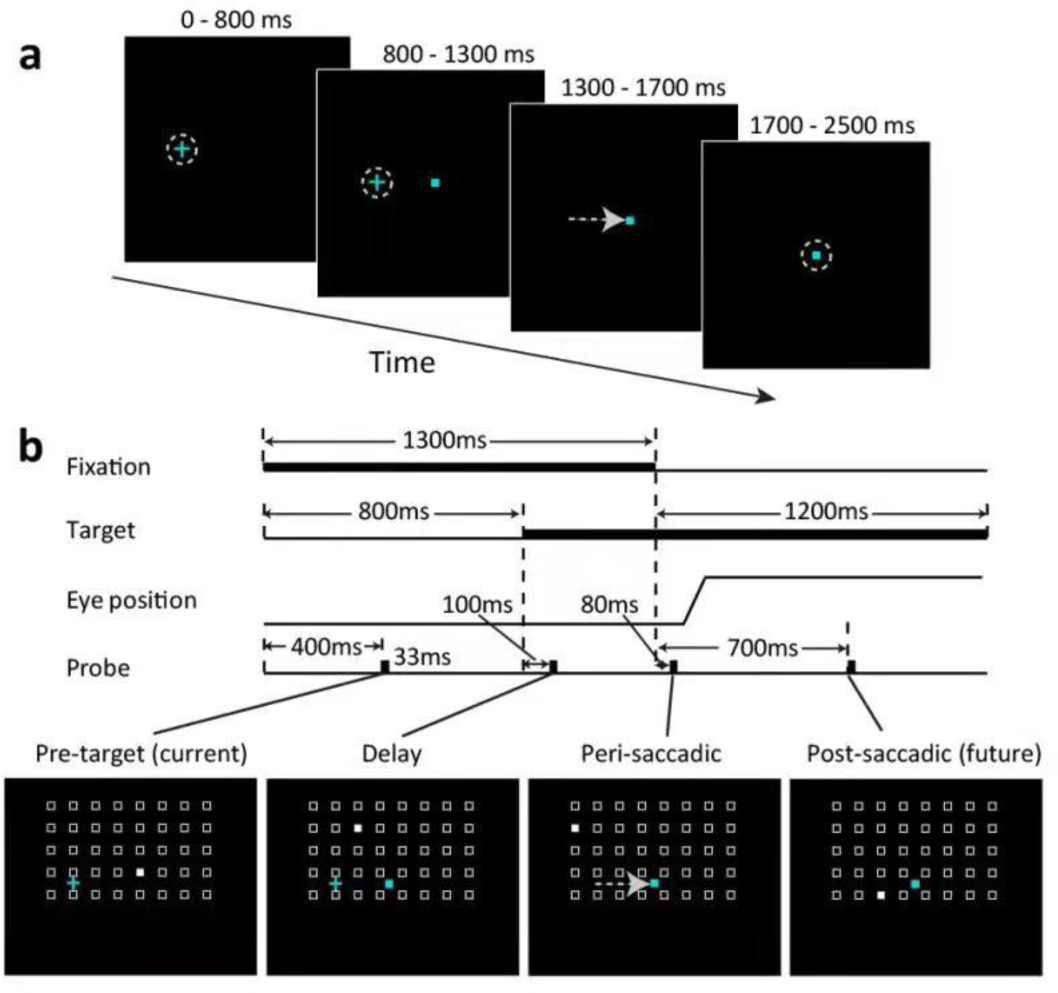
The delayed saccade paradigm. The small cyan cross and square represent the fixation point and the saccade target, respectively. (a) The trial sequence. The flashed probes are not shown here but see panel b. (b) The time courses of a trial. Four probes were flashed, one for each of the four epochs: pre-target (**c**urrent), **d**elay, **p**eri-saccadic, and post-saccadic (**f**uture). A cell’s RF mapped from these periods will be denoted **c**RF, **d**RF, **p**RF, and **f**RF, respectively. For each epoch, the probe stimulus (filled white square) appears randomly at one of the spatial array positions (open white squares, not shown in the experiment). The array size and location were tailored for each cell according pilot mapping.

### The direction and amplitude of RF remapping changed with time

We measured a cell’s remapping in the delay and perisaccadic periods as the shifts of its dRF and pRF centers relative to its cRF center, and defined the forward direction as the direction from the cRF center to the fRF center. We present the RF heat maps of example cells in Fig. 3 and the population shift directions in Fig 4. Only cells with significant RF shifts in the delay or perisaccadic periods (according to bootstrapping; see Methods) were included in the population analysis. For the delay period, the dRFs obtained 50 to 100 ms after the delay probe onset shifted slightly in directions between the initial fixation and the target (Fig. 3, second column; Fig. 4, first column), but 250 to 300 ms after the delay probe onset, the shifts turned more towards the target (Fig. 3, third column; Fig. 4, second column). For the perisaccadic period, the pRFs obtained 50 to 100 ms after the perisaccadic probe onset shifted towards the target (Fig. 3, fourth column; Fig. 4, third column). However, 25 to 75 ms after the saccade onset, the pRFs shifted mostly towards the fRFs (Fig. 3, fifth column; Fig. 4, fourth column). The mean shift directions were all significant except the early delay period of LIP (see the p values in Fig. 4 plots). We marked the 95% confidence intervals of the shift directions in Fig. 4. The mean directions changed significantly with time in both LIP and FEF (Fig. 4 caption). In Supplementary Figs. S1 to S5, we show that these results were robust against variations in analysis parameters.

**Fig. 3.**
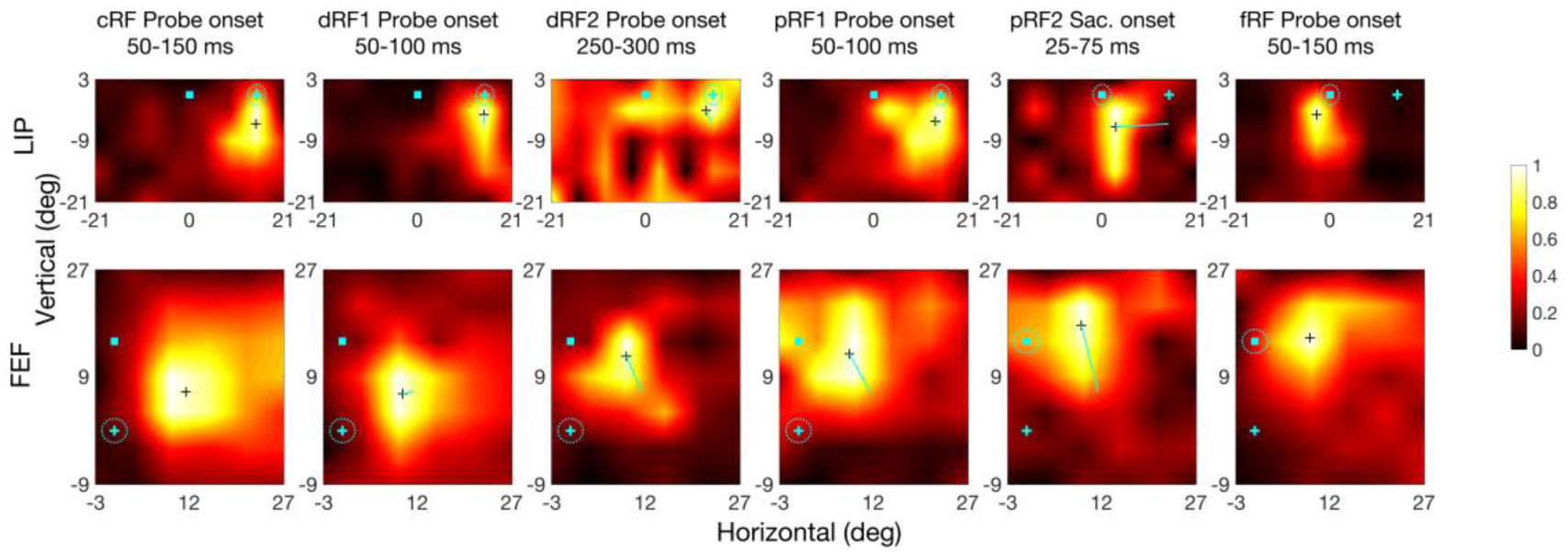
RF heat maps for an example LIP cell (top row) and FEF cell (bottom row) from different time periods (columns). In each heat map, the cyan cross and square indicate the initial fixation point and saccade target, respectively (see Fig. 2 for their time course), and the dashed cyan circle indicates the eye position. The small black cross in a map marks the RF center. The cyan lines in dRF and pRF maps indicate the center shift relative to the cRFs. The scale of normalized responses is shown on the right. The fifth column is based on saccade onset alignment of the repeated trials whereas the other columns are based on the probe onset alignment.

**Fig. 4.**
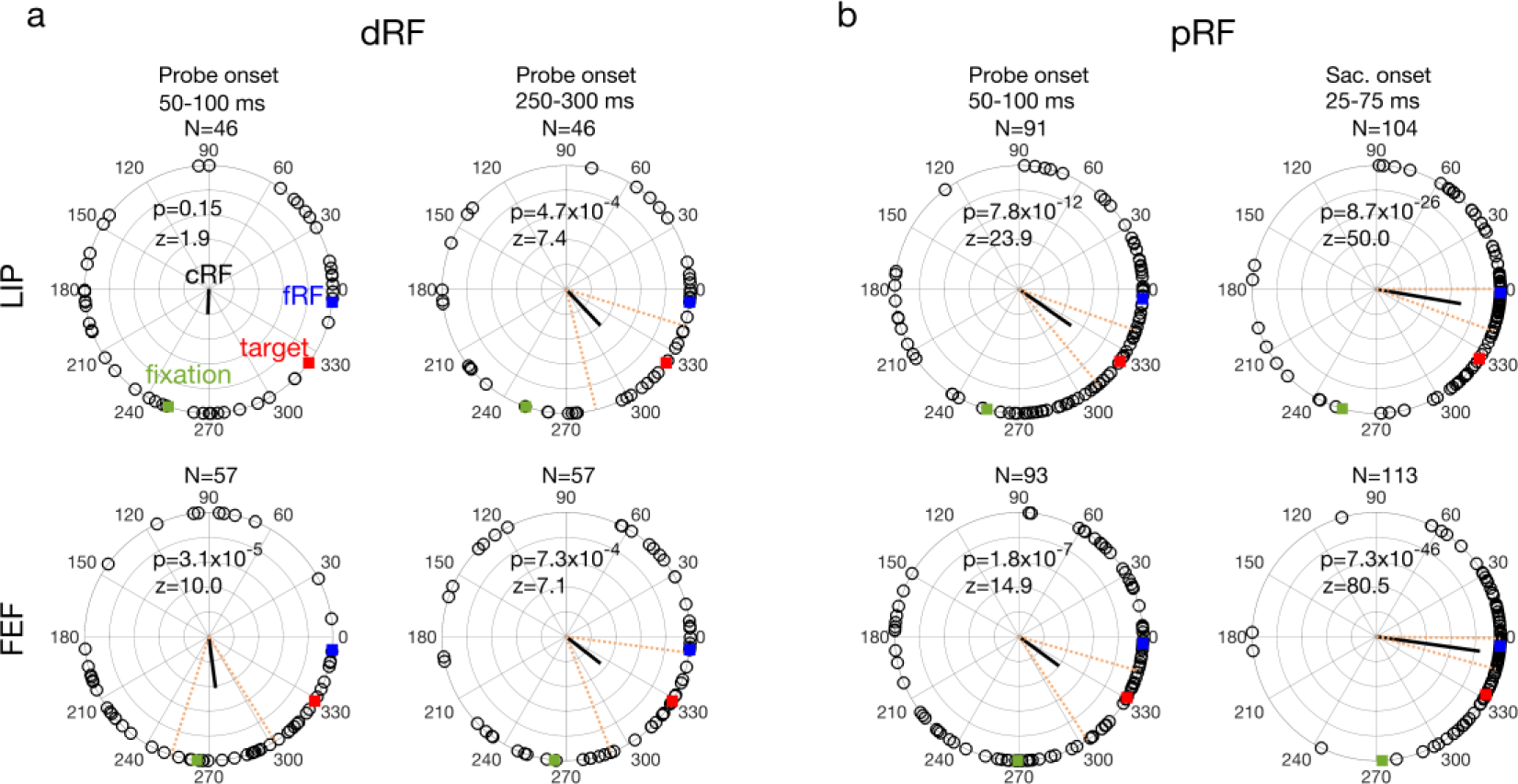
The delay (a: dRF) and perrisaccadic (b: pRF) shift directions of all LIP (top row) and FEF (bottom row) cells from different time periods (columns). In each polar plot, we align the cells’ cRFs at the center and saccade directions along positive horizontal. The cells’ mean fRF (forward), target, and initial-fixation directions are indicated by the blue, red, and green squares, respectively. Each open dot represents a cell’s RF shift direction. The thick black line represents the circular mean of all the shift directions, and its significance is indicated by the p values from Rayleigh test for circular distributions. The dashed red lines mark the 95% confidence interval of the mean direction. The mean directions changed significantly across time in both LIP (p = 1.0×10^−5^, F_3,283_ = 9.0) and FEF (p = 2.1×10^−9^, F_3,316_ = 15.5), with Watson-Williams multi-sample test. The fourth column is based on saccade onset alignment of the repeated trials whereas the other columns are based on the probe onset alignment. Note that the numbers of cells (N) of the panels are different because the screening method was applied to each area, epoch, and alignment separately.

To examine the time course of the delay and perisaccadic RF remapping in detail, we used a moving window of 50 ms to analyze the dRF and pRF shifts as a function of time ^21^. For the delay period, the shift direction changed gradually from a direction between the initial fixation and target to the target (Fig. 5a, green). For the perisaccadic period, the shift direction changed gradually from the target to the fRF (Fig. 5b, green). When the shift directions pointed towards the initial fixation and subsequently toward the target the shift magnitudes were at most only 55% and 67% of the distance between the cRF and the target in LIP and FEF, respectively. When the shift directions pointed to the fRF the shift magnitudes approached the distance between the cRF and fRF (Fig. 5, purple). This shift would provide an accurate transsaccadic update of the retinal location of the probe stimulus which was flashed before the saccade. When we used the larger time interval used by Zirnsak et al. (50-350 ms after the onset of perisaccadic probe), we did find the partial shifts toward the target that they described (Fig. 6a). This is not surprising given that the time interval they used included both the time in which we found convergent remapping and the time in which we found forward remapping. Note that the cells’ pRF centers were significantly closer to their fRF centers than to the targets (see tests in Fig. 6b), indicating that on average, the forward remapping was stronger than the convergent remapping during the perisaccadic period in both LIP and FEF.

**Fig. 5.**
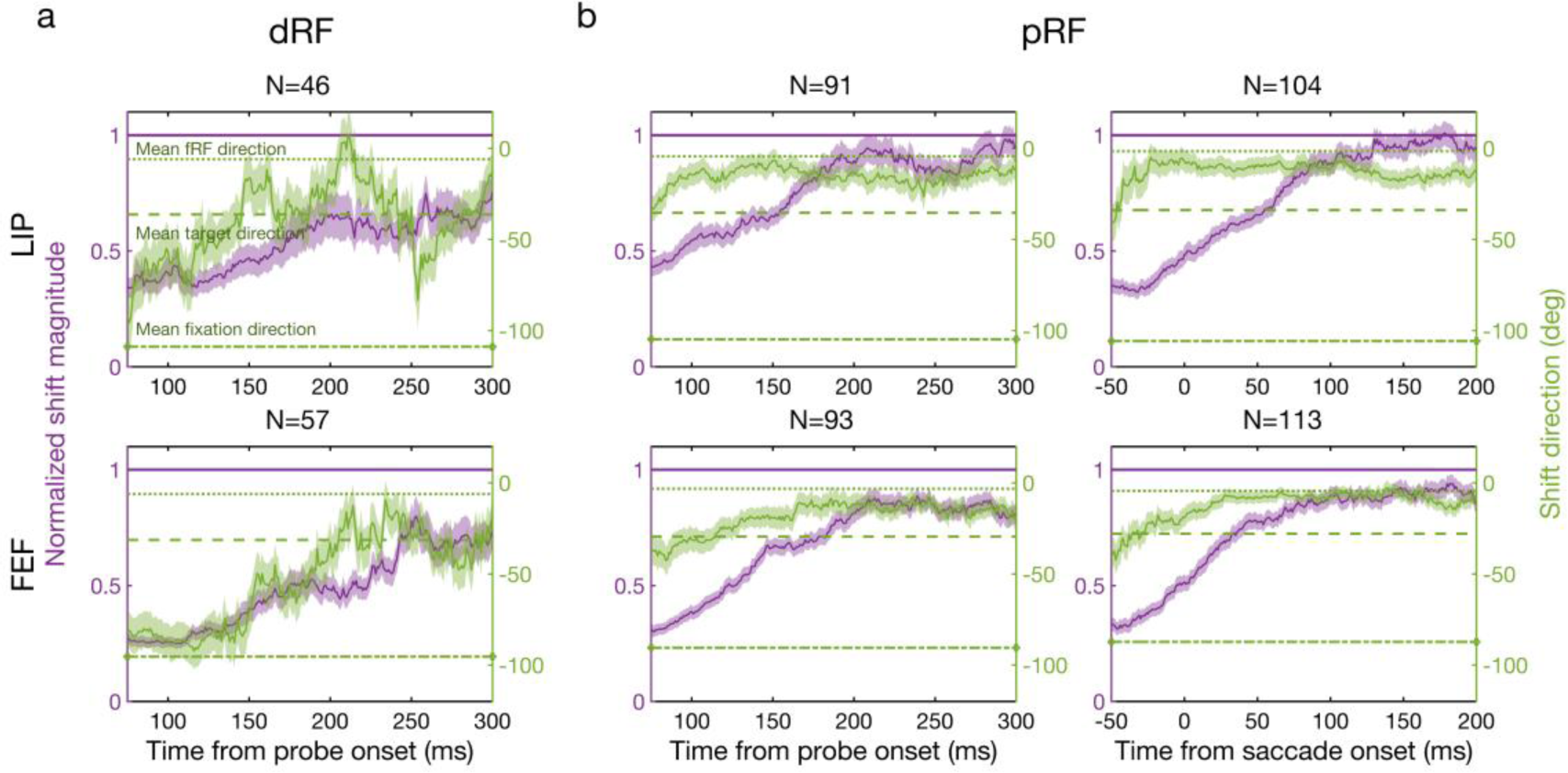
Remapping time courses of the delay (a: dRF) and perisaccadic (b: pRF) periods for LIP (top row) and FEF (bottom row). Each panel shows the mean normalized shift magnitude (purple curve, left y-axis) and the mean shift direction (green curve, right y-axis) as a function of time. The amplitude of one (purple horizontal line) means a shift magnitude equals the saccade magnitude. The average fRF, target, and initial-fixation directions are indicated by the green dotted, dashed, and dot-dashed lines, respectively. Each data point is calculated from the responses of the 50-ms window centered around that point. The light purple and green regions indicate 1SEM. The third column is based on saccade onset alignment of the repeated trials whereas the other columns are based on the probe onset alignment.

**Fig. 6.**
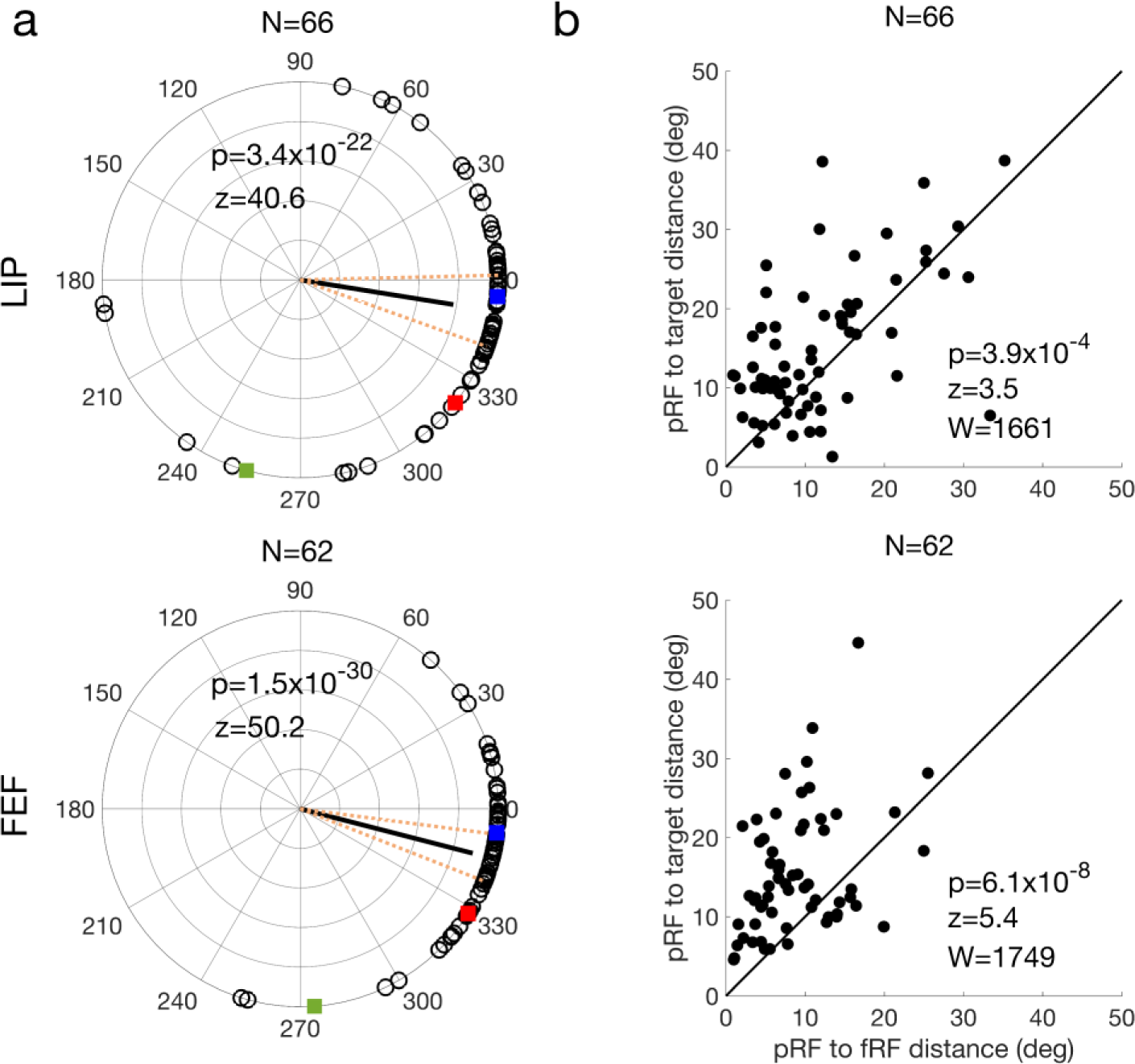
Analysis of our perisaccadic data using the method of Zirnsak et al. The LIP and FEF results are shown in the top and bottom rows, respectively. The polar plot of remapping directions (a) have the same format as that of Fig. 4. In (b), each cell’s pRF-to-target distance is plotted against its pRF-to-fRF distance. The p values in b indicate that for both LIP and FEF, the pRFs were significantly closer to the fRFs than to the targets on average (two-sided Wilcoxon signed-rank test).

### A circuit model for the forward and convergent remapping

To account for the data, we modeled a 2D array of cells with their RF centers topographically arranged (Fig. 7, black circles). A rightward saccade is to be made from the cross to the square, and we record from the filled black cell. The above physiological results suggest that two mechanisms may be responsible for the convergent and forward RF shifts, respectively. The *first* mechanism is an attention-modulated circuit for convergent shift. Inspired by the physiological evidence of center-excitatory/surround-inhibitory modulation of visual responses around the saccade target in both LIP ^19^ and FEF ^20^, we hypothesized center-excitatory/surround-inhibitory connections among all cells (Fig. 7, red lines; only the connections from the cell at the target location are shown). When there is attention at a location, we assume that connections from the cells tuned to that location are enhanced. This center/surround connectivity in the spatial domain is similar to that in the orientation domain ^22–25^. Such connectivity simulated convergent shifts of orientation tuning curves after perceptual learning or adaptation ^24–26^; here we used it to account for convergent shifts of spatial tuning (i.e., RFs) induced by attention.

**Fig. 7.**
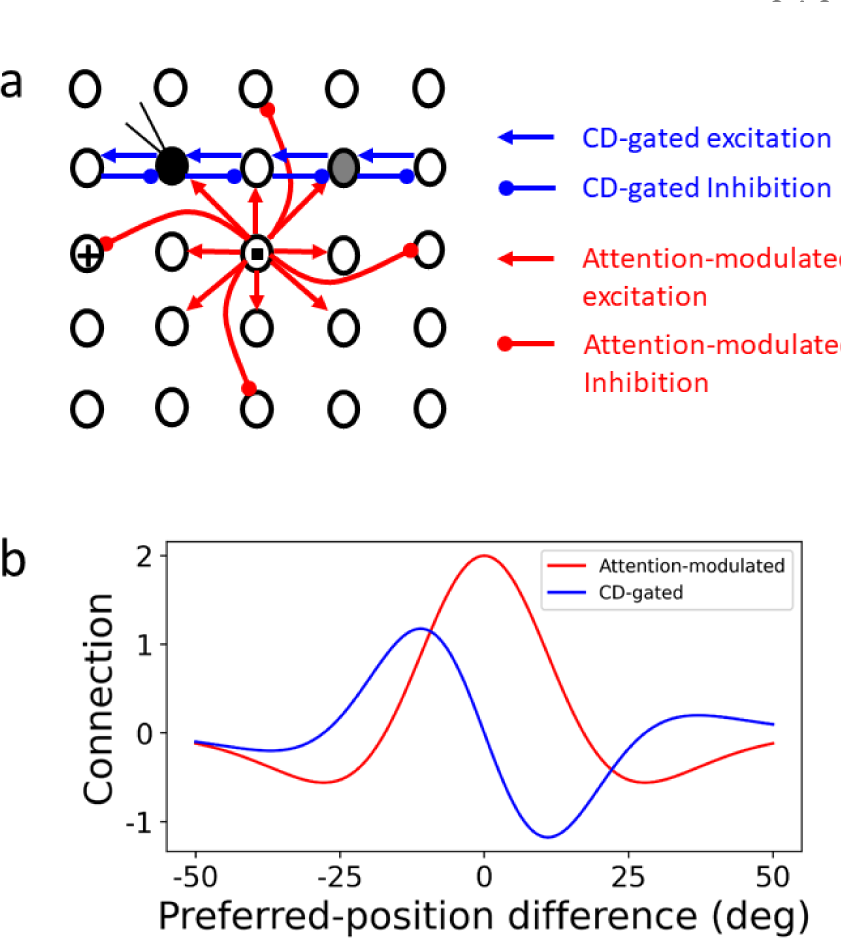
The circuit model for explaining both forward and convergent RF shifts. (a) Schematic model structure. Black circles represent RF centers of a 2D array of topographically arranged LIP or FEF cells. The small cross and square indicate the initial fixation and target positions, respectively. The filled black circle is the cRF of the cell under recording, and the gray circle is its fRF. Only a small fraction of the connections is shown for clarity. (b) The attention-modulated symmetric connectivity (red) and CD-gated anti-symmetric connectivity (blue, for rightward saccades) as a function of the difference between two units’ preferred locations (RF centers).

The *second* mechanism is the CD-gated directional connections that can explain perisaccadic forward expansion of RFs, as shown by Wang et al ^11^. The cells have CD-gated connections in all directions but for clarity, only a small subset of the connections for the second-row cells gated by the CD for a rightward saccade are shown (Fig. 7, blue lines). These connections are normally off. However, around the onset of the saccade, they are tuned on by the CD signals, allowing stimulation at the cell’s fRF (gray circle), and at the region between the fRF and cRF, to propagate to the recorded cell, generating perisaccaidc forward RF shifts.

We considered a 2D array of 50×50 LIP/FEF units covering a space of 50°×50°, each receiving feedforward visual inputs and recurrent inputs from other units via the attention-modulated center/surround connections and CD-gated directional connections. For the delay period, we divided the attention between the fixation point and the target, and in the perisaccadic period, we introduced the CD signal (Methods). We probed the model with flashes in the four epochs as in the experiment to measure cRFs, dRFs, pRFs, and fRFs. By weighting the two mechanisms differently, it is straightforward to generate cells with various degrees of forward and convergent shifts as we found in LIP and FEF (an example shown in Fig. 8).

**Fig. 8.**
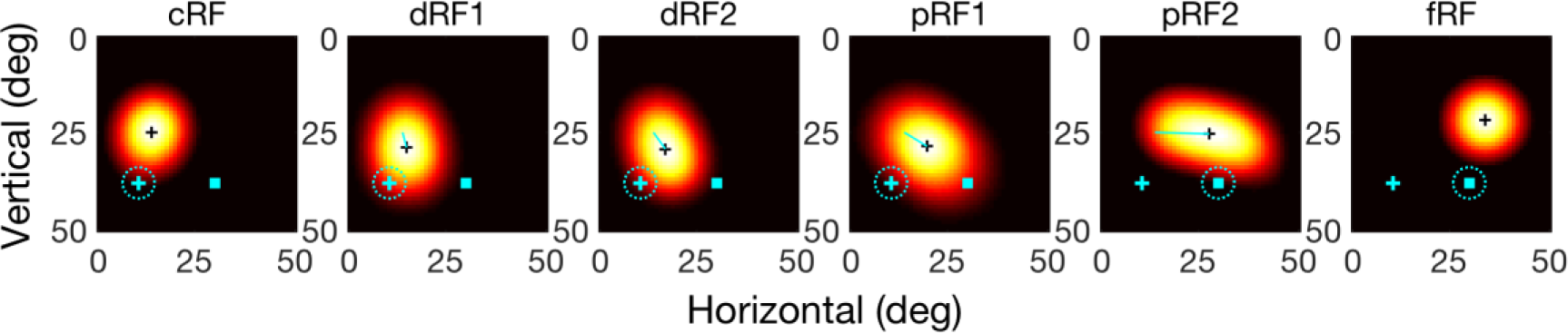
Simulations of RF shifts in the delayed saccade task of Fig. 2. The first and last panels represent a model cell’s cRF and fRF, respectively. The second and third panels represent the cell’s early (dRF1) and late (dRF2) RFs in the delay period. The fourth and fifth panels represent the cell’s early (pRF1) and late (pRF2) RFs in the perisaccadic period. In each panel, the cyan cross and square are the initial fixation point and target, respectively, and the dashed cyan circle indicates the eye position. The black cross marks the RF center. The thin cyan line in each dRF or pRF panel indicates the shift from the cRF center to the dRF or pRF center.

The model makes two predictions (Fig. 9, top row). 1) Since the CD-gated connections propagate neuronal responses from a cell’s fRF to its cRF, a distance equal to saccade amplitude, the pRF forward shift amplitude should increase with the saccade amplitude. 2) In contrast, the strengths of the attention-modulated center/surround connections depend on the distance from the attentional locus, so the pRF convergent shift amplitude should vary with the distance between a cell’s cRF center and the target. cRFs near the target have little room to shift toward the target and those far away are barely affected by attention at the target. There is thus an intermediate, optimal distance for maximal convergent shift. To test these predictions, we pooled the LIP and FEF cells whose pRFs from 0 to 100 ms after the saccade onset shifted between the fRF and target directions, and did a parallelogram decomposition of each shift vector into its forward and convergent components. The results (Fig. 9, second row) are consistent with the predictions. The scarcity of data at large cRF-to-target distance was due to technical limitations: when this distance is large, it was difficult to keep a cell’s entire cRF and fRF within the screen boundaries.

**Fig. 9.**
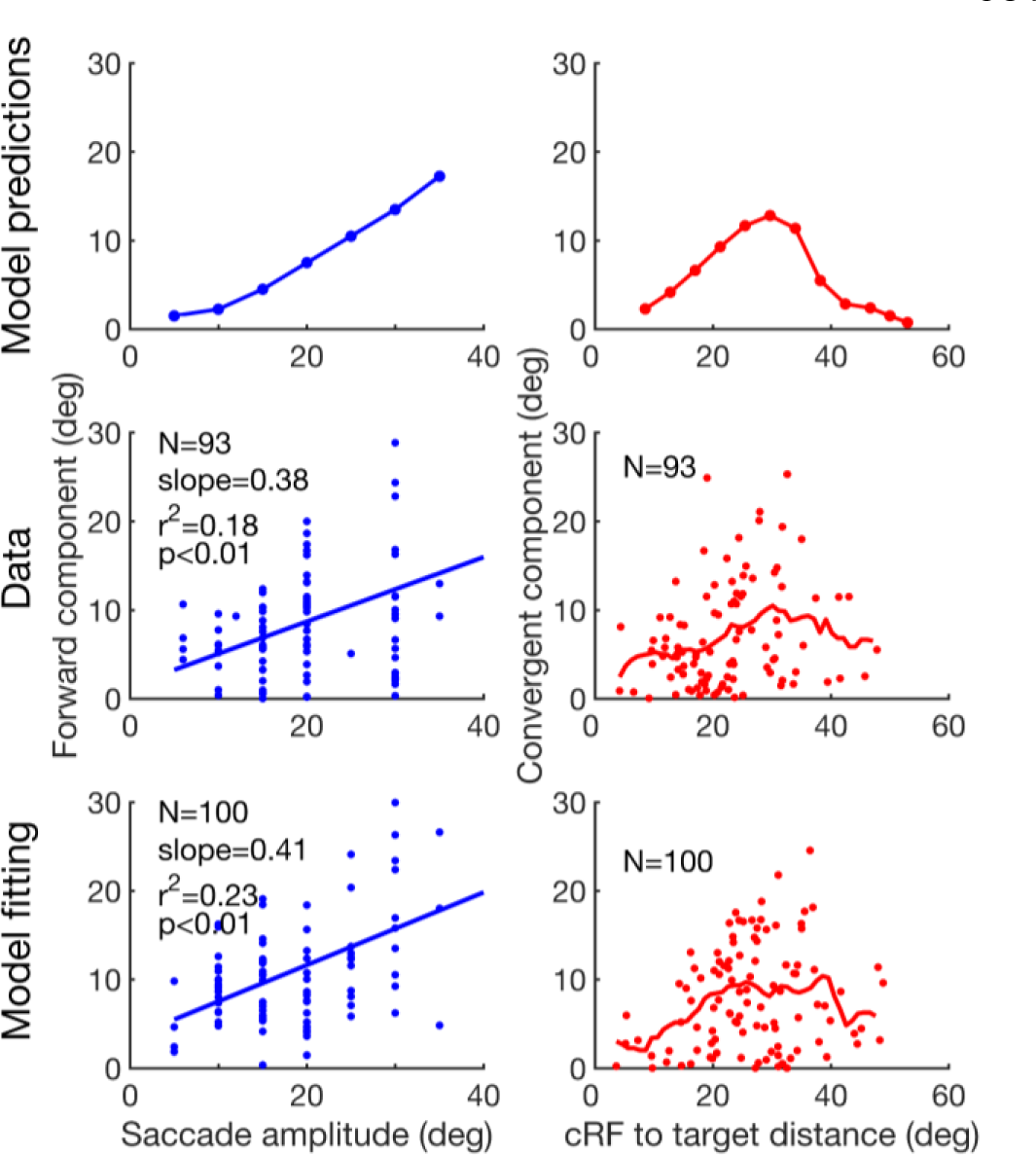
Test the model predictions. First row: the predicted forward shift amplitude as a function of saccade amplitude, and the convergent shift amplitude as a function of the cRF-to-target distance. Second row: the pooled LIP/FEF pRF data confirming the predictions. Third row: Fitting the model to the data. For the second and third rows, the blue lines in the left column are linear fits, and the red curves in the right column are the moving average with a window size of 10 deg. For the forward component, the slopes of the data and the model are not significantly different from each other (p = 0.79, t_189_ = 0.27, t-test). For the convergent component, the distributions of the data and the model are not significantly different from each other (p = 0.056, Peacock’s 2D Kolmogorov-Smirnov test).

To explain our data more quantitatively, we obtained distributions of the model parameters by fitting the model to the perisaccadic LIP and FEF data, and interpolated each parameter distribution as a mixture of Gaussians (Methods). We then randomly resampled parameters from these distributions and run the model to obtain the pRF shifts. The results (Fig. 9, bottom row) matched the data well.

### The emergence of the required connectivity patterns in neural networks trained to update retinal positions across saccades

Both the attention-modulated center/surround connections and the CD-gated directional connections in our circuit model are motivated by physiological evidence in FEF and LIP. Nevertheless, one may argue that the model is ad hoc, designed specifically to explain the forward and convergent remapping. Is there a simple, functional consideration that leads to both connectivity patterns automatically? As we noted above, pRF remapping appears to update the retinotopic location of remembered (and disappeared) stimuli across saccades, a requirement for performing the double-step memory saccade task (Introduction). We therefore hypothesized that the two sets of connections in the circuit model are for such transsaccadic updating, with the center/surround connectivity for storing the retinal position of a stimulus of interest ^27,28^, and the CD-gated connectivity for updating the memory across saccades ^6,11,29^.

We tested this hypothesis by training neural networks on the predictive updating task and examining whether randomly initialized connections converge to the required patterns after training. For simplicity, we considered horizontal saccades only. The neural networks consisted of two layers of units. The first layer provided visual inputs to the second layer which simulated LIP/FEF cells. The second-layer units were trained to produce activity patterns representing the correct retinal locations of input stimuli across saccades without reafference delay. The connections from the first- to second-layer units were convolutional so that retinotopic inputs specified the feedforward component of LIP/FEF RFs. The second-layer units were fully and recurrently connected to each other with three sets of weights. The first two sets were gated by CD signals for saccades of opposite directions whereas the third set was not gated by the CD signals but could be modulated by attention (Methods). All connection weights in the network were randomly initialized. The visual input in the first layer was a Gaussian bump centered at the initial retinal location of a stimulus (Fig. 10a, left, for a 50 ms stimulus). Two additional input units provided CD signals for opposite saccade directions. The desired output in the second layer was the same Gaussian bump centered at the correct retinal position of the (disappeared) stimulus across saccades (Fig. 10a, middle, for a rightward saccade started at 150 ms and thus a leftward displacement of the representation for the stimulus’ retinal position). The weights were trained by minimizing the quadratic difference between the actual and desired outputs. Many variations of the simulation produced similar results (see Supplementary Information).

**Fig. 10.**
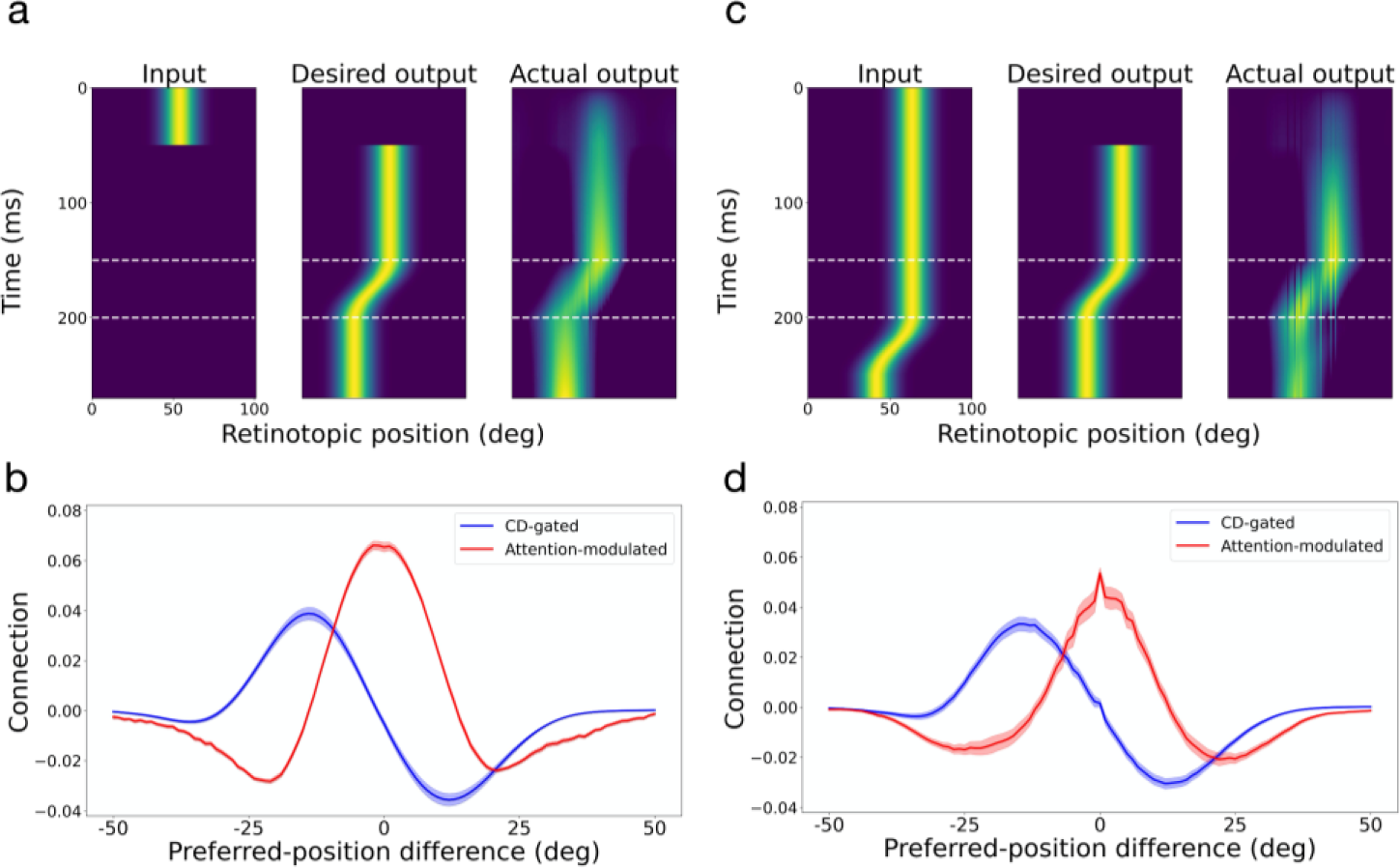
Automatic generation of the required connectivity patterns in the circuit model by training neural networks to predictively update retinal positions of stimuli across saccades. Panels a and b are for the case of brief input stimuli and panels c and d are for the case of persistent input stimuli. Panels a and c show the test input (not included in the training), the desired output, and the actual output. Panels b and d show the average connection weights as a function of the difference between two units’ preferred positions (RF centers). The red and blue curves are for the attention-modulated and CD-gated connections, respectively. Only the CD-gated connections for rightward saccades are shown. The shaded areas indicate 1SEM.

After the training converged and the actual output resembled the desired output for test inputs not used during the training (Fig. 10a, right), we determined the units’ mean connection weights to other units as a function of the distance between their preferred positions. Fig. 10b shows the results, with the attention-modulated and CD-gated connections in red and blue, respectively. The CD-gated connections shown are for rightward saccades and the mirror pattern for leftward saccades was also learned (not shown). Remarkably, these connectivity patterns closely resemble those we chose for the circuit model in Fig. 7b. Note that the connections in Figs. 7b and 10b are comparable only in their shapes, but not in their scales. This is because the circuit model and the artificial neural networks used different scales to represent the units’ activities.

We then repeated the above neural network training but with persistent visual stimuli (Fig. 10c). Interestingly, we obtained very similar results, with not only the CD-gated directional connections but also the center/surround connections (Fig. 10d). The reason is that during saccades, the desired output position is different from the delayed reafference input position (Fig. 10c, middle and left). The symmetric center/surround connectivity is needed to create an attractor activity pattern which was updated by the asymmetric CD-gated directional connectivity, independent of the input activity pattern^30^. We also considered the case where both brief and persistent input stimuli were trained together and where the attentional modulation was turned off, and again obtained similar results (see Supplementary Figs. S7 and S8).

We conclude that the connectivity patterns required by the circuit model emerge automatically and robustly in neural networks trained to update the representation of stimuli’s retinal positions across saccades. This result suggests that although the center/surround connections and the CD-gated connections in the circuit model are for explaining convergent and forward RF shifts, respectively, they work synergistically to enable transaccadic stability.

## Discussions

In the 19th century, Herman von Helmholtz, not only a great physicist but also a pioneering ophthalmologist, examined a patient who was blind in one eye from diabetes and sustained a paralysis of the lateral rectus muscle ^5^. When the patient tried to look in the direction of the paralyzed muscle he perceived that the visual world moved in the opposite direction and then drifted back. Helmholtz postulated that under normal conditions, the brain uses the oculomotor signal to feed back to the visual system and adjust for the saccade. The discovery of perisaccadic forward remapping ^6–10^ provided a physiological mechanism for Helmholtz’s theory. Neuropsycholgical evidence also supports this theory: patients with parietal lesions ^31,32^ cannot perform the double-step task when the subject makes the first saccade in the direction contralateral to the lesioned cortex. Furthermore, monkeys with inactivation of the medial dorsal nucleus of the thalamus, which relays CDs of saccadic commands from the SC to the FEF, cannot perform the double-step task, and their FEF neurons do not exhibit perisaccadic forward remapping ^29,33^. However, an influential study questioned the existence of forward remapping ^15^ and instead showed that when they analyzed the activity of FEF neurons in the interval from 50 to 350 ms after the appearance of probe, which appeared around the time of the saccade target, many neurons seemed to remap their receptive fields toward the target.

In humans ^34^ and monkeys ^17^ the abrupt onset of a visual stimulus evokes attention as measured by a change in perceptual threshold, as does the planning of a saccade ^17,35^. Because of the large time interval (50-350 ms after probe onset) used by Zirnsak et al., we wondered if their results might be confounded by the presence both of an attentional event (the appearance of the saccade target) and the generation of a saccade, which would result in the combination of convergent remapping evoked by the saccade target and forward remapping evoked by the CD of the saccade command. Here, we addressed this controversy by using a delayed saccade task to separate the appearance of the target and the generation of the saccade. We recorded from LIP and FEF with matched procedures, and found that LIP and FEF showed similar patterns of remapping which varied with time: during the perisaccadic period, RFs converged toward the target shortly after the probe onset, but around the time of saccade and onward, the shift directions became predominantly forward toward fRF. When we used a large time window to integrate perisaccadic activities ^15^, the pRFs were closer to the fRFs than to the targets, indicating stronger forward remapping than convergent remapping. We further found that the convergent shift started in the delay period when the saccade command, and its CD, must be suppressed; and the shift direction turned from between the initial fixation and the target to the target. Thus, unlike the forward shift that depends on the saccade CD ^29^, the convergent shift appeared to follow attention from the initial fixation to the target. We conclude that both types of remapping are present in FEF and LIP and that forward remapping is not an artifact of undersampling convergent remapping. The convergent and forward RF shifts may be viewed as attentional remapping and perisaccadic remapping, respectively. These two types of remapping have also been found in V4 ^21^ but with a much slower time course than that in LIP/FEF, raising the possibility that V4 might inherit the remapping from LIP/FEF.

Because our delayed-saccade paradigm helps distinguish between the forward- and convergent-remapping mechanisms, we were able to construct a circuit model for both types of remapping. Specifically, we integrated attention-modulated center/surround connections and CD-gated directional connections to explain the convergent and forward RF shifts, respectively. The model’s predictions on the forward shift amplitude as a function of the saccade amplitude and the convergent shift amplitude as a function of the cRF-to-target distance agreed with the data. We then showed that both sets of connections emerged automatically and robustly in neural networks trained to update representations of retinal positions across saccades. Since this updating is needed for the double-step memory saccade task, it can be viewed as an operational definition of transaccadic stability. We suggest that the CD-gated connections and the center/surround connections together specify a mechanism for transaccadic stability. The mechanism follows a classic prescription ^30^: symmetric center/surround connections produce attractor dynamics to represent a stimulus as an activity bump whereas the asymmetric CD-gated connections move the activity bump for updating across saccades. Although we initially used the center/surround connections to explain convergent remapping, they might be an integral part of transaccadic stability mechanism.

Zirnsak et al ^15^ suggested that convergent RF remapping explains compressive perceptual mislocalization: stimuli flashed briefly before a saccade are perceived as occurring at the saccade target when postsaccadic visual references are present ^36,37^. However, whether convergent remapping produces compressive mislocalization is unclear, and depends on, among other things, whether the positional decoder is aware of the remapping ^38^. Additionally, when saccadic adaptation is used to dissociate the postsaccadic eye position and the target position, the perceived compression is toward the eye position, not the target position ^39^ whereas LIP and SC neurons represent the target position, not the eye position ^40,41^. Therefore, one would not expect convergent remapping (in LIP and SC at least) to explain compressive mislocalization.

In addition to perisaccadic RF remapping, a prominent physiological finding relevant to transaccadic perceptual stability is gain fields, the modulation of visual response by eye position ^42^. Whereas perisaccadic remapping may realize the stability by predictively updating retinal representations across saccades, gain fields may do so by combining eye position and retinal representations to form head-centered representations ^43^. Recent studies suggest that gain fields and perisaccadic remapping may be responsible for transaccadic stability at long and short time scales, respectively ^44,45^, consistent with their respective dependence on slow proprioceptive eye-position signals and fast saccade CD signals ^29,46^. On the other hand, it is theoretically possible to integrate fast CD signals to provide fast, predictive eye-position signals. The existence of such CD integrators is an open question for future research.

Our study may also have functional implications on potential relationships between working memory and attention. According to our models, although the center/surround connections may store working memories of visual stimuli including saccade targets, the same connections can be modulated by attention to generate convergent RF shifts. The relationship between working memory and attention has been discussed in the literature ^47^. Our work, however, suggests a specific mechanism: attention to a stimulus modulates the connections that are responsible for storing the stimulus in working memory. Therefore, LIP and FEF circuits might integrate mechanisms for working memory, attention, saccade planning, and transaccadic stability all together.

## Methods

### Animal preparation

Three male adult rhesus monkeys (*Macaca mulatta*) weighing from 9 to 11 kilograms participated in this study. All procedures were approved by the ethics committee at Beijing Normal University. We have complied with all relevant ethical regulations. We surgically planted two search coils (one for each eye; Crist Instrument Sclera, sample rate at 2.7KHz), a head restraint post, and two recording chambers (for LIP and FEF, respectively; PEEK), for each monkey. We positioned the recording chambers according to our experience and/or the MRI scans. We centered the LIP chambers for the three monkeys, respectively, at 3, 10, and 3.2 mm posterior to the interaural plane, and 13, 15, and 15 mm lateral from the middle line. We centered the FEF chambers for the three monkeys at 28, 18, and 23.5 mm anterior to the interaural plane, and 13, 15, and 18 mm lateral from the middle line. The two recording areas were verified later (see below).

### Recording procedures

We used insulated tungsten microelectrodes (0.3∼1.0 MΩ, FHC) to record single-unit activity. We inserted the electrodes though dura via stainless steel guide tubes, and controlled their advancement in the cortices with micromanipulators (NAN Instruments). Neuronal activities collected by the electrodes were amplified (Alpha Omega) and filtered (268-8036Hz) before online sorting with AlphaLab SnR (Alpha Omega). We identified LIP based on persistent activities in the delay period of a memory saccade task ^48^, and FEF according to micro-stimulation (100ms, 0.05mA, biphasic pulses) evoked saccades of fixed vectors ^49,50^. After several recording sessions, we did MRI scan of the first two monkeys’ LIP chambers and the third monkey’s LIP and FEF chambers, confirming that the LIP recording sites were within the lateral bank of the intraparietal sulcus, and the FEF recording sites were in the anterior bank of arcuate sulcus. The same recording and analysis procedures were applied to LIP and FEF.

After isolating a single unit with a template-matching method, we first did a pilot mapping of its visual RF: while the monkey maintained central fixation in each trial, we flashed a sequence of 6 probe stimuli (1°×1°) at random locations sampled from an 8×8 array with adjacent locations separated by 6° in both horizontal and vertical directions. A probe lasted 33 ms and successive probes were separated by 400 ms. Each location had about five responses. If visual inspection determined that the unit showed clear responses for at least one probe location, we moved on to the main, delayed saccade task (Fig. 2). We tailored the array of probe positions to cover the unit’s cRF-fRF region and the target region, according to the pilot RF mapping and the planned saccade target for the unit. Across cells, the array varied from 4×5 to 10×12 positions, with 5×8, 5×9, and 6×8 the most common. The spacing between adjacent positions (along both horizontal and vertical axes) varied from 2° to 6°, with 6° the most common. The saccade amplitude varied from 5° to 30°, with 15° and 20° the most common. Despite our effort, the RFs of some cells were not measured sufficiently complete because of the limited screen size and large RFs and/or large saccades; these cells were excluded (see below). As shown in Fig. 2 and described in the text, the delayed saccade task allowed us to measure a cell’s cRF, dRF, pRF, and fRF from the initial-fixation (**c**urrent), **d**elay, **p**erisaccadic, and postsaccadic (**f**uture) epochs of a trial.

In the actual experiments, the initial fixation point and the target were both red squares of 0.3° width, but for the ease of illustration, we represented them as cyan squares and crosses, respectively, in the figures.

### Data screening and analysis

We screened and processed the data as follows. (1) We selected the cells with significant visual responses. For each epoch, we aligned the repeated trials to the probe onset. For the perisaccadic epoch, we additionally aligned repeated trials to the saccade onset. For each epoch and probe position of a cell, we calculated the response as the mean firing rate 50-150 ms after the probe onset or 0-100 ms after the saccade onset (these windows were chosen because they contained most of the activities), and the baseline as the mean firing rate 0 to 50 ms before the probe onset. We found the probe position that had maximal response, and then performed a single Wilcoxon rank-sum test (two-sided) against the corresponding baseline at the 0.05 level. This procedure avoided multiple comparisons. Cells were selected separately for each epoch and alignment. For a selected cell in an epoch, we followed Zirnsak et al ^15^ to normalize its spatial responses according to (r_k_-r_min_)/(r_max_-r_min_) for all k, where r_k_ is the response at position k, and r_max_ and r_min_ are the maximum and minimum responses across all positions. An advantage of this normalization is that because of the subtraction of r_min_, any non-visual (such as saccade related) responses are discounted. Also note that because we always place the target outside a cell’s RF, saccade contribution to the measured responses must be minimal. (2) We selected cells with well-measured RFs. For each epoch and alignment of a cell, we linearly interpolated the normalized responses across positions to obtain the RF heat map. We traced the response contour at 85% of the maximum (contour criterion) and required that 80% of contour were within the sampled position grid (completeness criterion). We then estimated the RF center as the center-of-mass of the responses within the region set by the contour criterion. We used the 85% contour criterion instead of Zirnsak et al.’s ^15^ 75% because the higher value determined the RF center more reliably. (In Supplementary Information Figs. S1 to S5, we demonstrate that changing the two criteria do not change our conclusions.) (3) We selected cells with significant RF shifts. For each cell, we calculated the shifts of its dRF and pRF centers relative to its cRF center in visual angles, and determined the significance of a shift via the following bootstrapping ^51^. For each epoch and probe position of a cell, we assumed that the spike count of a trial followed a Poisson distribution with the mean equal to the measured mean spike count. We then simulated the recording and analysis of the cell 1000 times by sampling spike counts from the distributions with the trial numbers equal to those of the actual experiment. To determine, for example, whether the 1000 dRF centers shifted significantly from the 1000 cRF centers in the simulation, we calculated their overlaps along the axis linking the mean dRF center and cRF center, and required the overlap to be less than 5%. After the screening, we investigated how remapping changed with time by choosing various 50 ms windows to determine dRFs and pRFs as detailed in the main text.

We considered each recorded cell as a distinct sample. After the screening (see the paragraph above), the numbers of cells for different time periods were different, and for this reason, results from different time periods could not be treated as repeated measures. We thus used Watson-Williams test to determine whether remapping directions changed significantly over time in Figs. 4 and S1-S5.

We also applied Zirnsak et al’s method to analyze perisaccadic remapping in our LIP and FEF data. In addition to screening for cells with sufficient trials, well-sampled RFs, and significant RF shifts in perisaccadic epoch, we selected the trials in which perisaccadic probes occurred within 150 ms before saccade onset, for all epochs used the responses from 50 to 350 ms after the probe onset, and set the contour criterion to 75% ^15^.

We used Matlab to perform the data screening and analysis.

### Circuit model

We simulated a 2D array of 50×50 LIP/FEF units covering a space of 50°×50°, each unit governed by the standard equations:

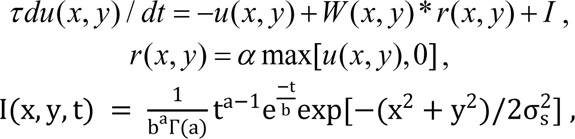

where *u*(*x*, *y*) and *r*(*x*, *y*) are, respectively, the membrane potential (relative to spike threshold) and firing rate (relative to background rate) of the unit at location (*x*, *y*), *τ* = 20 ms is the membrane time constant, *α* is a constant relating *u*(*x*, *y*) to *r*(*x*, *y*) (which affects the model only through its product with *W*, specified below), * denotes spatial convolution, *W* specifies connections between the units, and *I* represents the feedforward visual inputs with *a* = 2, *b* = 18 ms. For each unit, its connection matrix *W* (*x*, *y*) to other units at relative coordinates (x, y) is a sum of two parts: (1) center-surround connections modeled as a difference between two circularly symmetric 2D Gaussians:

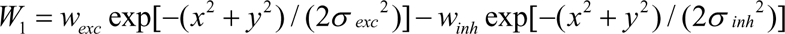

(with *α w_exc_* = 4, *α w_inh_* = 2, *σ _exc_* = 12°, *σ_inh_* = 18°), and (2) asymmetric connections with excitation in the opposite direction of the pending saccade ^11^; for horizontal saccades, we used the antisymmetric form: *W*_2_ = *β* ∂ *W*_1_ / ∂*x* along the horizontal axis, where β = 12 determines the relative strength between *W*_1_ and *W*_2_ . (For other saccade axis, *W*_2_ should be rotated to the saccade axis.) Many expressions for *W*_2_ would work; we choose the derivative of *W*_1_ for its known property in shifting activity profile in the direction where *W*_2_ is excitatory ^30^ and for reducing the number of free parameters. Note that Wang et al^11^ used the CD-gated excitatory connections against the saccade direction and a global inhibition. We used an equivalent connectivity with inhibitory and excitatory connections along and against the saccade direction, respectively. Fig. 7b shows the shapes of *W*_1_ and *W*_2_ along *x* for rightward saccade. For the delayed saccade task, *W*_1_ is multiplied by an attentional modulation factor:1+ *w_att_* exp[−(*x*^2^ + *y*^2^) / (2*σ _att_* ^2^)] centered at the attentional locus (initial fixation or target position), where *σ _att_* = 15° . Similarly, *W*_2_ is multiplied by a CD gating factor: *w_CD_* exp[−(*t* / *σ*)^6^ / 2] where *w_CD_* is not independent but determines the CD strength through its product with β, *t* is measured relative to the saccade onset time, and *σ _CD_* = 65 ms. For the six time periods in Fig. 8 (cRF, dRF1, dRF2, pRF1, pRF2, and fRF), we set *w_att_* at the initial fixation to 0.4, 0.8, 0.6, 0, 0, and 0, *w_att_* at the target to 0, 0.2, 0.3, 0.45, 0.2, and 0, and *w_CD_* to 0, 0, 0, 0.1, 0.6, and 0, respectively. *w_att_* at the initial fixation was larger for dRF1 and dRF2 than for cRF because after the target onset, the monkeys had to suppress any saccades to the target and maintain the initial fixation. Visual inputs from a stimulus to the LIP/FEF units were modeled as a circular Gaussian centered at the stimulus with *σ _s_* = 7° . We probed the model with flashes in the four epochs as in the experiment to measure cRFs, dRFs, pRFs, and fRFs. Many variations of the model and/or the parameters produced similar results. For example, the attentional modulation and CD-gating functions can be replaced by simple step functions, and the parameters can be optimized to fit individual cell’s RF shifts (see below). We implemented the circuit model with COSIVINA, an open source toolbox for Matlab.

The model predicts that the forward-shift amplitude grows with the saccade amplitude and that the convergent-shift amplitude depends on the cRF-to-target distance, with a maximal shift at an intermediate distance (see text). To make these predictions more quantitative, we obtained distributions of the model parameter set Θ = (*w_exc_, σ_exc_, w_inh_, σ_inh_, w_att_, σ_att_, w_CD_*) by fitting the model to the perisaccadic LIP and FEF data. We focused on these parameters as they are most relevant to the RF shifts. We first initialized the parameters to random values, each drawn from the uniform distribution *U* (0, 20) . For each recorded cell with a shift vector 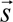, we used PyTorch’s differentiation engine and Adam optimizer (both learning rate and weight decay set to 0.01) to perform gradient descent on the parameters by minimizing the cost function:

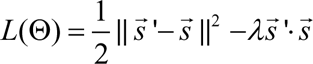

where 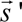 is the shift vector produced by the model. The first term minimizes the difference between 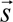 and 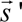. We included the second, dot-product term because when 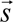 and 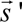 have small amplitudes, the first term can be small even when 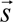 and 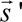 point at different directions. The second term ensures that the two vectors point in similar directions. We let *λ* = 0.01as it produced good fits for all cells. Because of the limited number of recorded cells, we pooled together the optimized parameters obtained from fitting both the LIP and FEF cells, and for each saccade amplitude, we fit each parameter distribution through the Gaussian kernel density estimation. We treated the parameters independently as we did not have nearly enough data to determine their joint distribution. We then sampled parameters from these distributions and run the model to predict the forward-shift amplitude as the function of the saccade amplitude, and the convergent-shift amplitude as a function of the cRF-to-target distance (Fig. 9). The cRF centers were resampled from the distribution of the measured cRF centers by fitting a mixture of Gaussians.

### Artificial neural networks

We trained artificial neural networks to predictively update retinal locations of stimuli across saccades and demonstrated automatic and robust emergence of both the center/surround connections and the CD-gated directional connections needed in the circuit model for explaining the convergent and forward RF shifts, respectively. A network consisted of two layers of units: the first layer provided visual inputs, originated from retina, to the second, LIP/FEF layer. For simplicity, we only considered the horizontal dimension and horizontal saccades. Each layer had 100 units representing 100° of space. The connections from the first to second layer was translationally invariant ^52^ (convolution kernel size of 5°) so that the second layer preserved the retinotopic representation of the first layer. There were two additional input units with a one-hot representation of the CD signals for the two opposite directions of saccades. For a given saccade, the relevant CD unit was turned on for the duration of the saccade. The second-layer units are fully and recurrently connected with three sets of weights. The first two sets were multiplicatively gated by the two CD input units for opposite saccade directions, respectively, whereas the third set was optionally modulated by attention when the stimulus was considered task relevant (e.g., when it was the saccade target). The simulations in Figs. 10 and S7 included attentional modulation with *σ_att_* = 15° and *w_att_* = 0.4. In Fig. S8, we showed an example where we obtained similar connectivity patterns in the absence of attentional modulation. The dynamics of the units was governed by equations identical to those for the circuit model above, and we used the ReLU activation function.

The networks were trained on the task of predictively updating the retinal position of visual stimuli across saccades. Specifically, the output units should have the activity pattern representing the correct retinal position of an input stimulus across saccades without reafference delay. Both the input and desired output are Gaussian activity patterns with *σ* = 6° . We considered both brief and persistent input stimuli, with one stimulus per training trial. The brief stimuli appeared for 50 ms before saccades and then disappeared whereas the persistent stimuli stayed for the duration of the simulations.

Importantly, the input units provided inputs to LIP/FEF units and we assumed a 50 ms delay from retina to LIP/FEF (Fig. 10c, left). The output activity pattern was trained to compensate for this delay by using the CD signals (Fig. 10c, middle). Therefore, regardless of whether the input stimuli were brief or persistent, the desired output was the same: an activity pattern representing the correct retinal position of the stimuli without delay. This is what we mean by “predictive updating.”

The model was trained to minimize the mean squared error between desired output and actual output as follows:

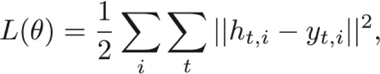

where *h_t,i_* and *y_t,i_* are the desired and actual outputs for unit *i* at time step *t*. All weights were randomly initialized by a uniform distribution 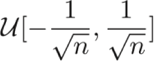, where *n* is the total number of weights of a given type (feedforward or recurrent) a unit receives. For the feedforward weights, *n* = 7 (the convolutional kernel size), and for the recurrent weights *n* = 100 (the number of recurrent units). We updated the weights with the Adam optimization algorithm (learning rate was 0.001, weight decay was 0.01). The model was implemented in PyTorch.

## Supplementary Information

Our physiological conclusions are robust against variations in the data analysis. There were two key criteria in our analysis. (1) Contour criterion: For a given RF heat map we measured, we set a percentage of the peak response to mark a contour around the peak for calculating the RF center of mass. The contour criterion was set to 85% in the main text. (2) Completeness criterion: We required that the measured RF heat map included a minimum percentage of the contour defined by the contour criterion. This completeness criterion was set to 80% in the main text. We did extensive additional data analysis to demonstrate that our physiological conclusions are robust against variations in these criteria. We focused on Fig. 4 of the main text as it contained the main results on the RF shift directions in the delay and perisaccadic periods for both LIP and FEF. In Figs. S1 and S2, we kept the contour criterion at 85%, but set the completeness criterion to 90% and 70%, respectively (instead of 80% in Fig. 4). In Figs. S3 to S5, we changed the contour criterion to 75%, a value used by Zirnsak et al.’s ^15^, and set the completeness criterion to 90%, 80%, and 70%, respectively. These figures all show results similar to those in Fig. 4 of the main text.

**Fig. S1.**
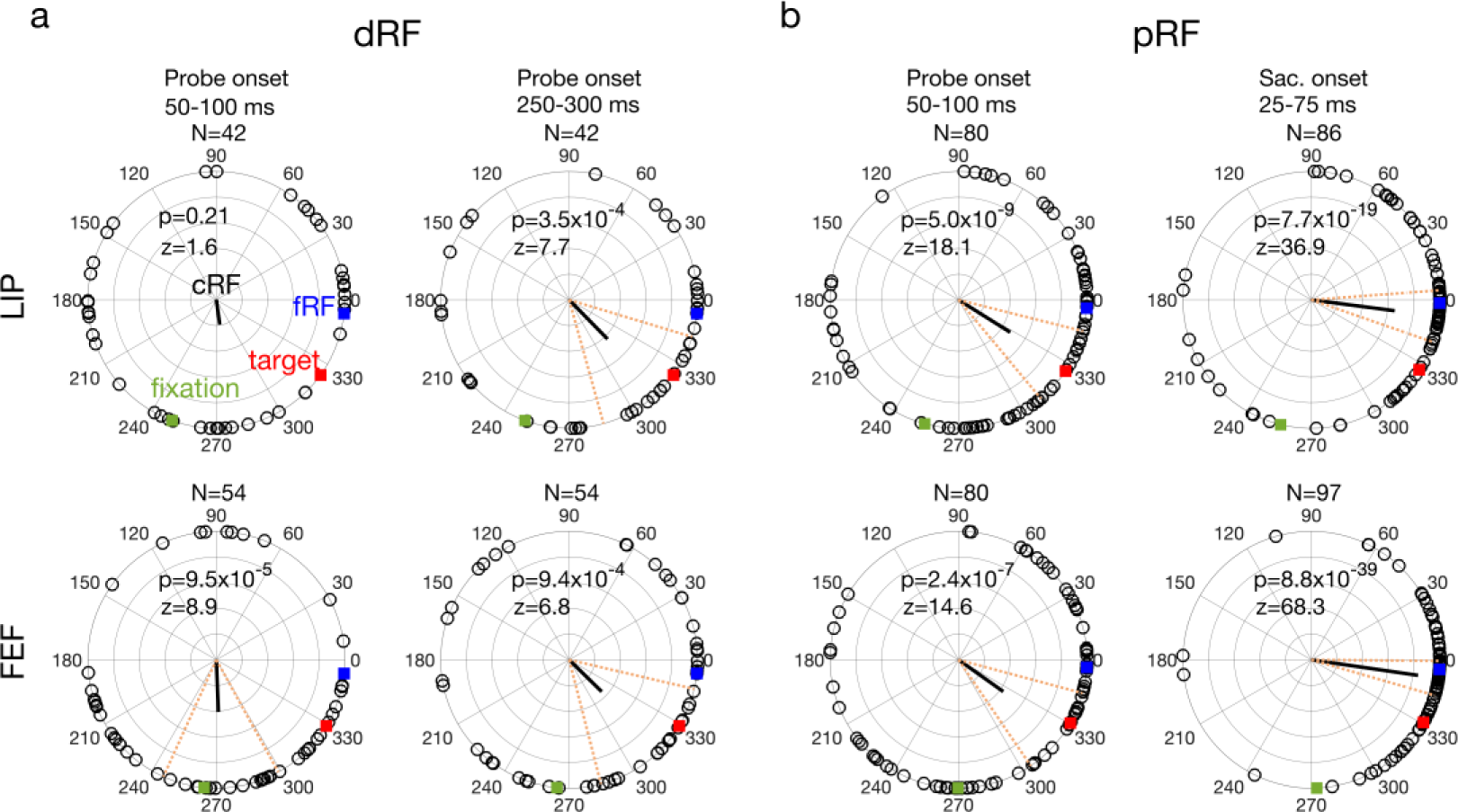
The delay (a: dRF) and perrisaccadic (b: pRF) shift directions of all LIP (top row) and FEF (bottom row) cells from different time periods (columns). The contour criterion was 85% and the completeness criterion was 90%. The format was identical to that of Fig. 4 of the main text. The mean directions changed significantly across time in both LIP (p = 2.5×10^−4^, F_3,246_=6.6) and FEF (p = 1.9×10^−9^, F_3,281_=15.7), with Watson-Williams multi-sample test.

**Fig. S2.**
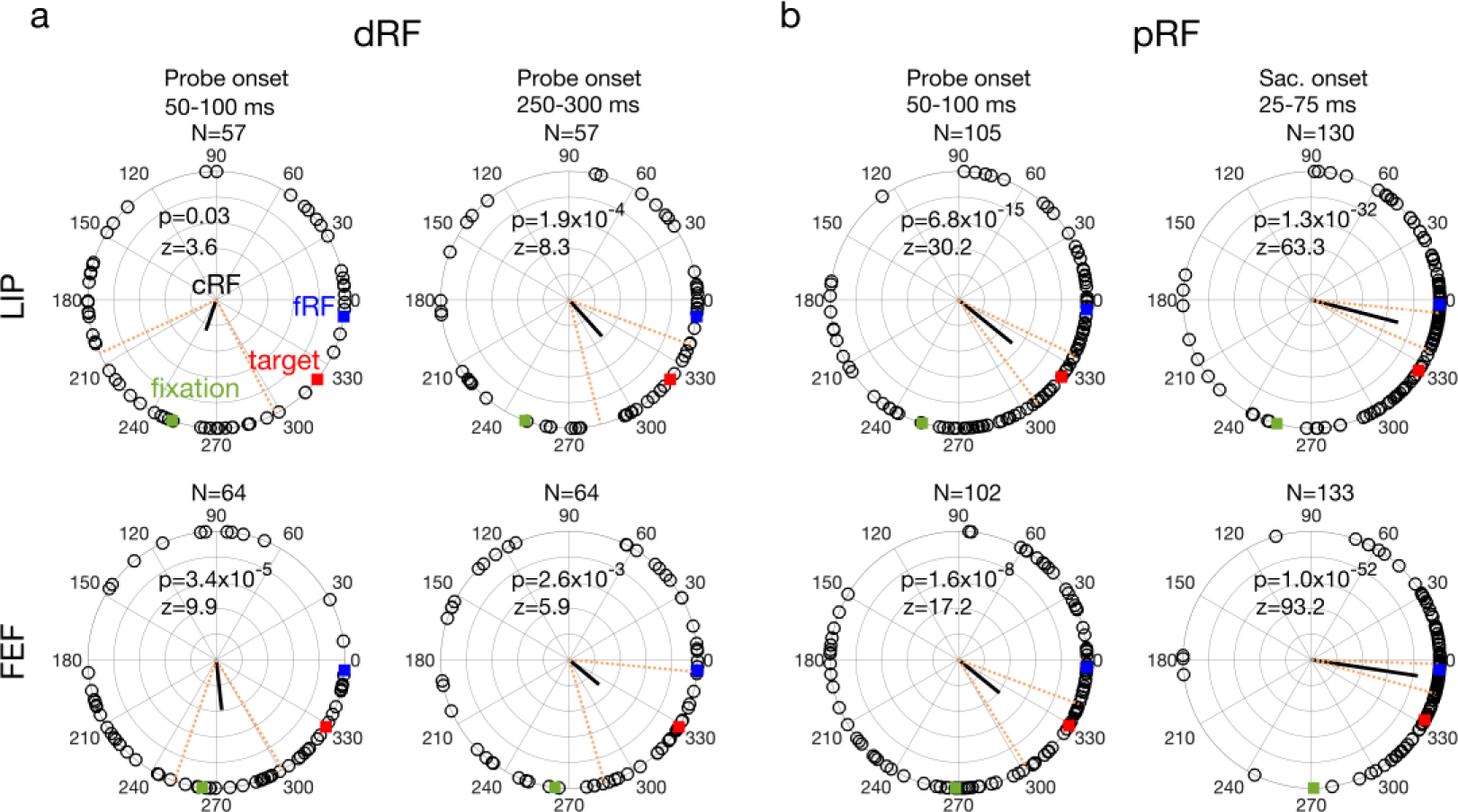
The delay (a: dRF) and perrisaccadic (b: pRF) shift directions of all LIP (top row) and FEF (bottom row) cells from different time periods (columns). The contour criterion was 85% and the completeness criterion was 70%. The format was identical to that of Fig. 4 in the main text. The mean directions changed significantly across time in both LIP (p = 3.6×10^−9^, F_3,345_=14.9) and FEF (p = 1.7×10^−10^, F_3,359_=17.3), with Watson-Williams multi-sample test.

**Fig. S3.**
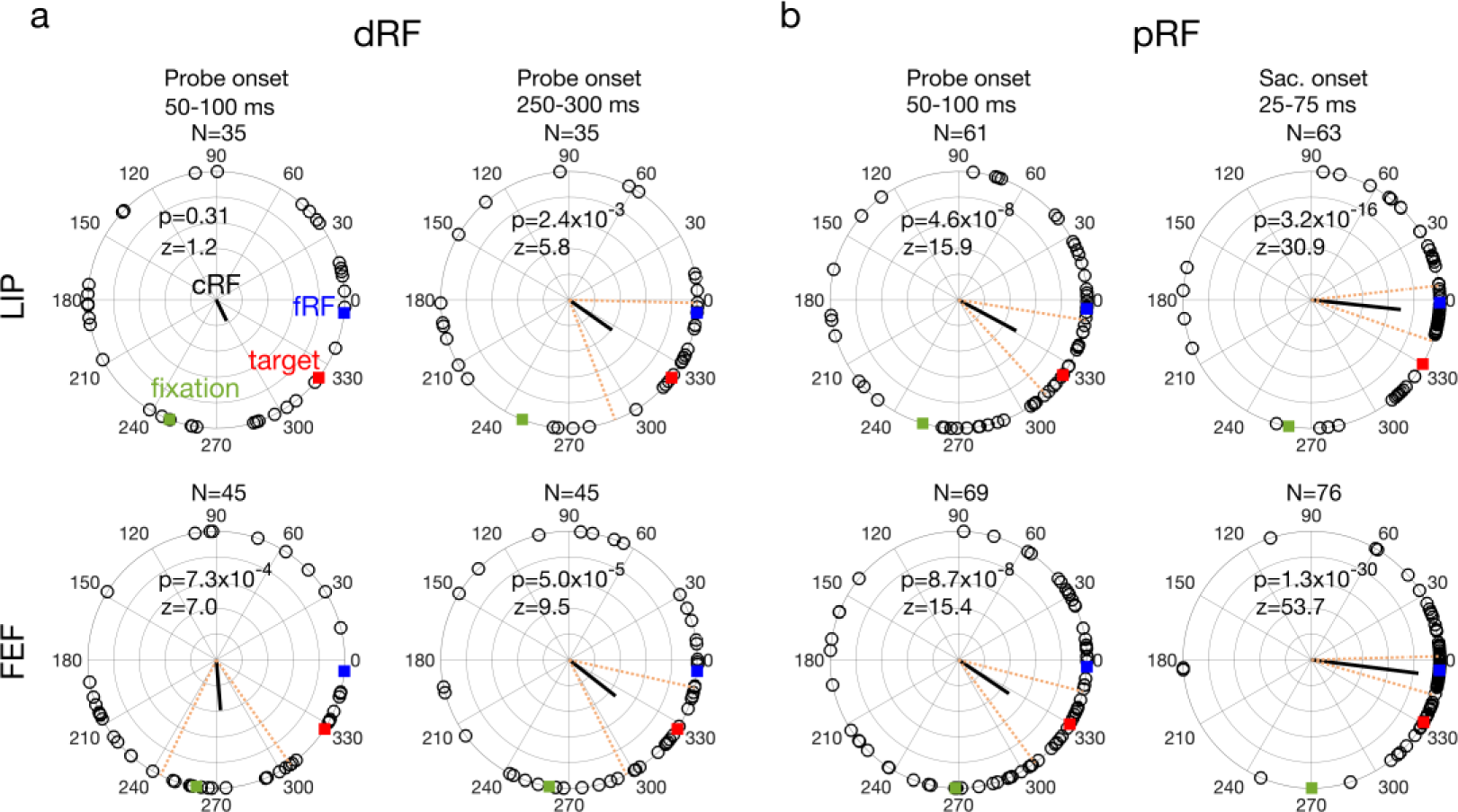
The delay (a: dRF) and perrisaccadic (b: pRF) shift directions of all LIP (top row) and FEF (bottom row) cells from different time periods (columns). The contour criterion was 75% and the completeness criterion was 90%. The format was identical to that of Fig. 4 in the main text. The mean directions changed significantly across time in both LIP (p = p = 0.021, F_3,190_=3.3) and FEF (p = 1.4×10^−7^, F_3,231_=12.5), with Watson-Williams multi-sample test.

**Fig. S4.**
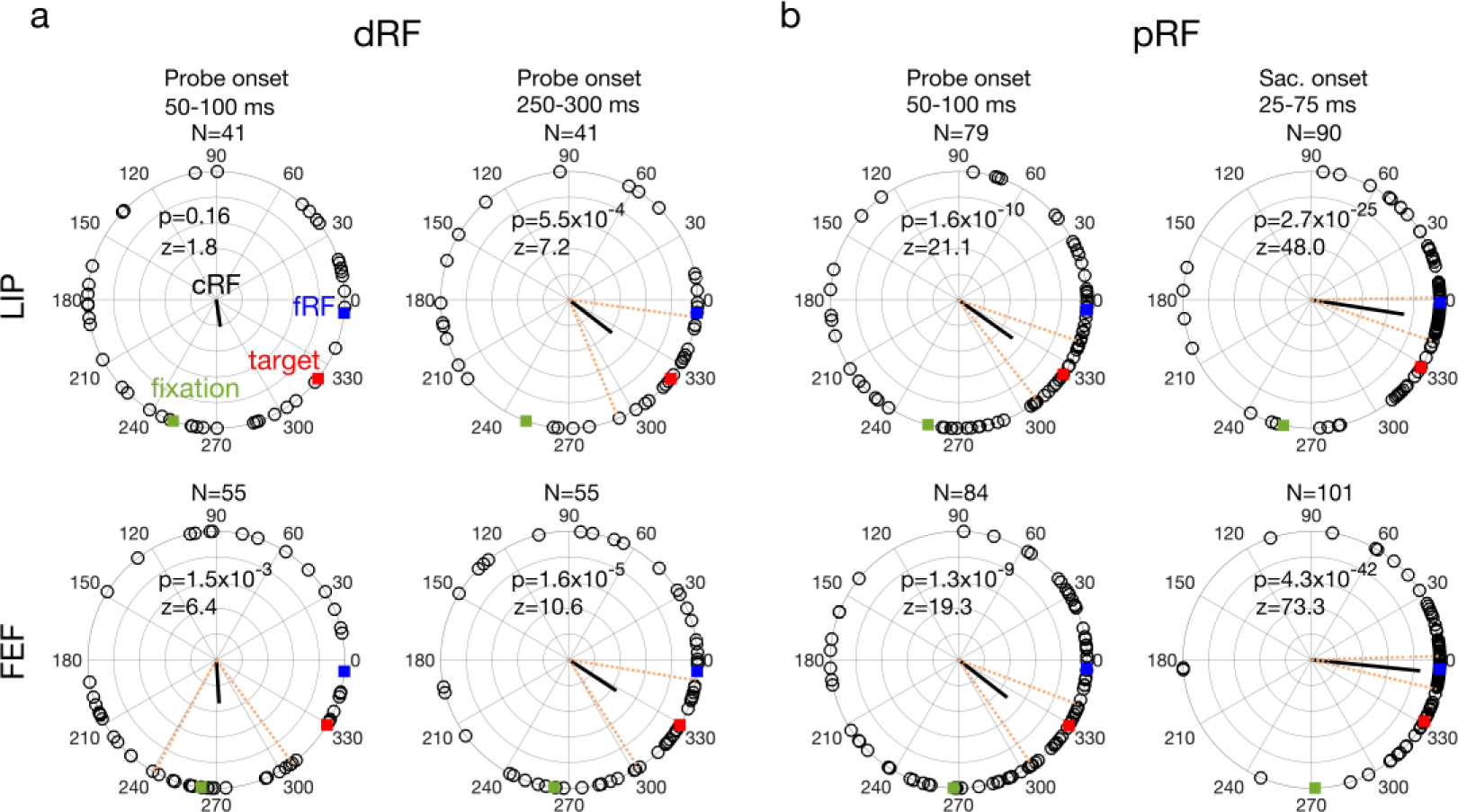
The delay (a: dRF) and perrisaccadic (b: pRF) shift directions of all LIP (top row) and FEF (bottom row) cells from different time periods (columns). The contour criterion was 75% and the completeness criterion was 80%. The format was identical to that of Fig. 4 in the main text. The mean directions changed significantly across time in both LIP (p = 1.9×10^−4^, F_3,247_=6.8) and FEF (p = 2.5×10^−9^, F_3,291_=15.4), with Watson-Williams multi-sample test.

**Fig. S5.**
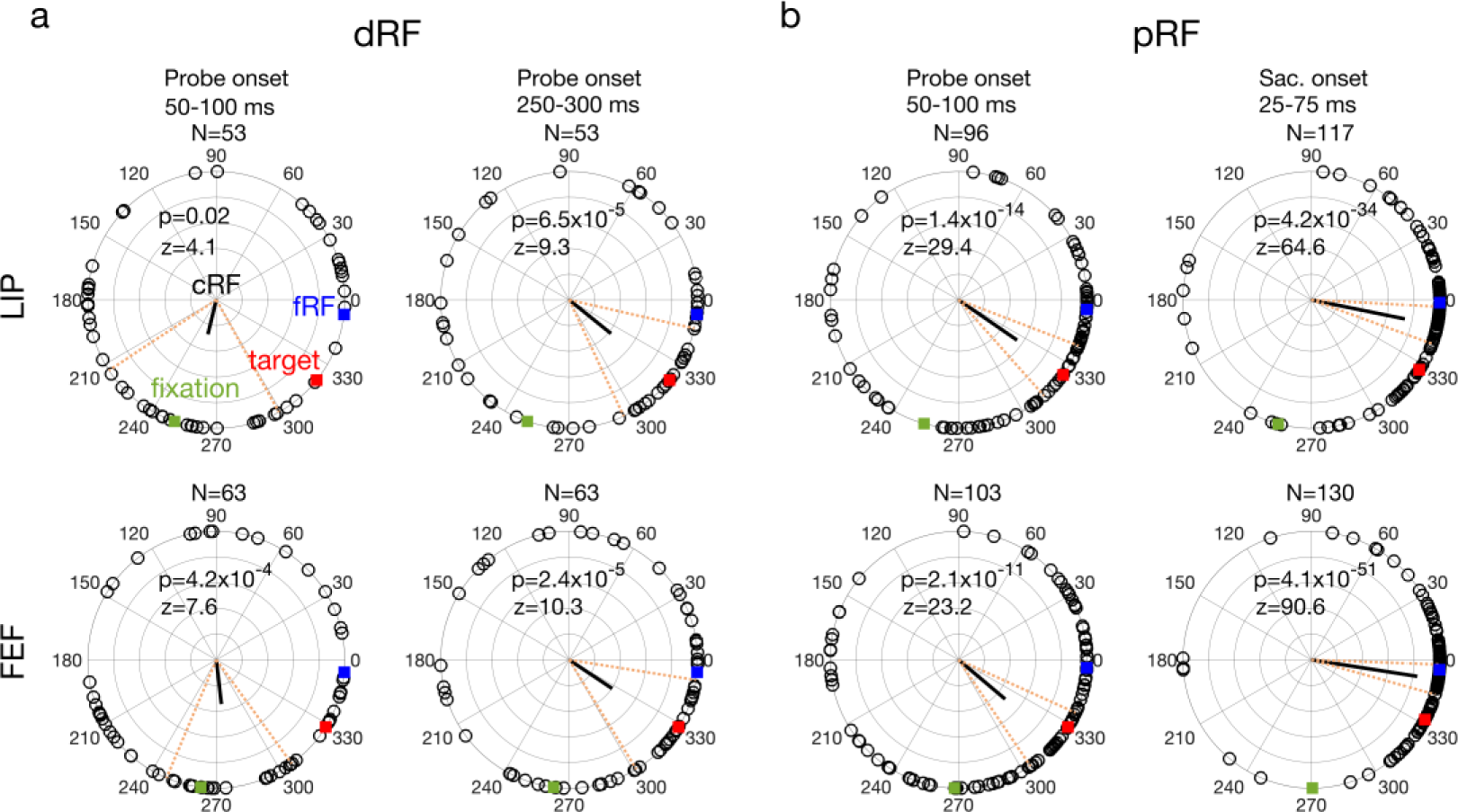
The delay (a: dRF) and perrisaccadic (b: pRF) shift directions of all LIP (top row) and FEF (bottom row) cells from different time periods (columns). The contour criterion was 75% and the completeness criterion was 70%. The format was identical to that of Fig. 4 in the main text. The mean directions changed significantly across time in both LIP (p = 6.7×10^−9^, F_3,315_=14.5) and FEF (p = 7.0×10^−10^, F_3,355_=16.2), with Watson-Williams multi-sample test.

In the main text, we showed the distributions of the RF shift directions at four time points (Fig. 4). For completeness, we show in Fig. S6 the distributions of the shift vectors (both the directions and amplitudes) ^21^.

**Fig. S6.**
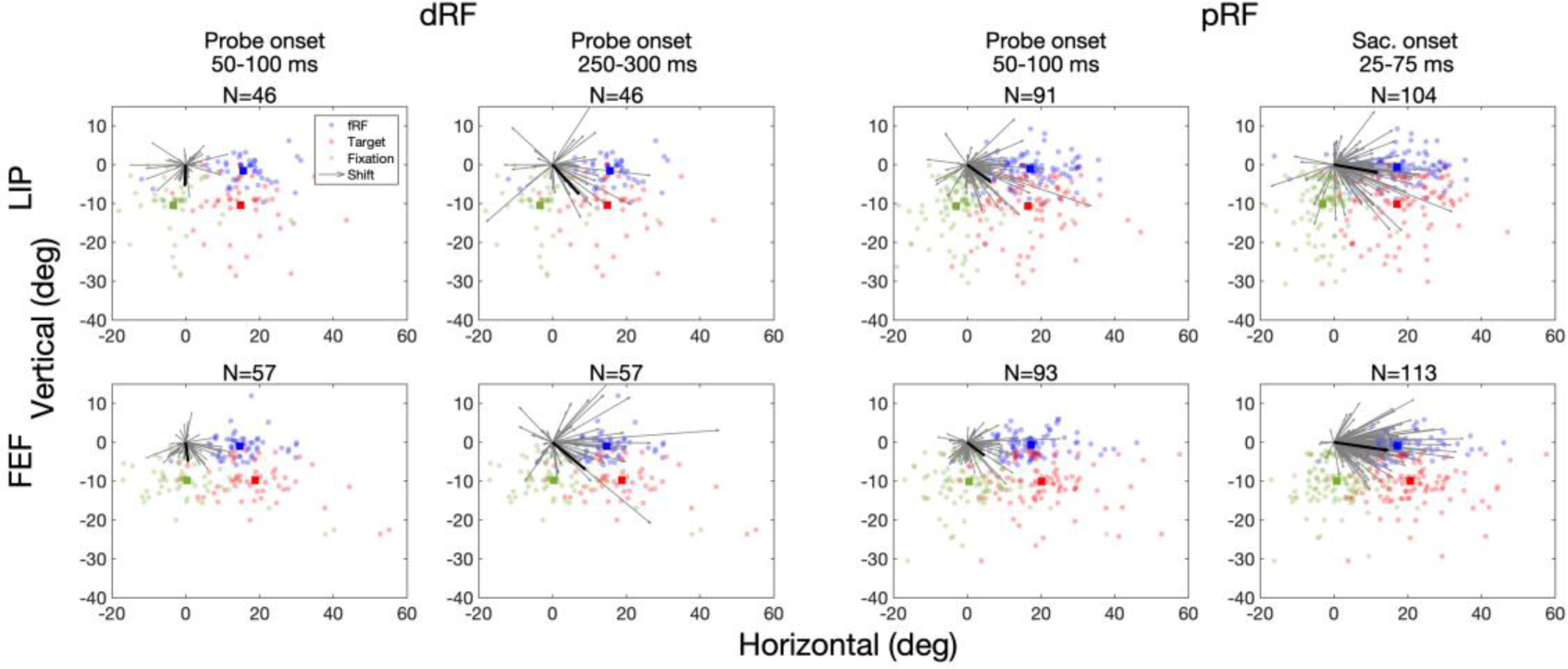
The delay (a: dRF) and perrisaccadic (b: pRF) shift vectors of all LIP (top row) and FEF (bottom row) cells from different time periods (columns). This figure corresponds to Fig. 4 of the main text but shows both the shift direction and amplitude of each cell. In each panel, we align the cells’ cRF centers at (0, 0) and saccade directions along positive horizontal. The cells’ fRF centers, the targets, and the initial-fixation points are shown as blue, red, and green dots, respectively, and their mean positions as the blue, red, and green squares, respectively. Gray arrows indicate the cells’ RF shift vectors and the black line is the vector determined by calculating the mean direction and mean amplitude of the individual vectors.

In Fig. 10 of the main text, we showed the automatic emergence of both the attention-modulated center/surround connections and the CD-gated directional connections in artificial neural networks trained to predictively update, across saccades, the representation of retinal locations of briefly flashed stimuli. We ran additional simulations to show that the same was true under many other conditions, with two examples in Figs. S7 and S8. In Fig. S7, we trained a neural network on both brief input stimuli and persistent input stimuli. In Fig. S8, we repeated the simulation in Fig. 10 of the main text but without the attentional modulation at the stimuli. In both cases, we found similar connectivity patterns to those in Fig. 10. It is not surprising that attention at the stimuli is not important for learning the connectivity patterns. To perform the task of updating the stimulus retinal positions, a network had to develop the center/surround connectivity to maintain the attractor activity pattern and the CD-gated directional connectivity to move the attractor pattern appropriately ^30^. These requirements do not depend on attentional modulation. Once the connections are learned, attention can modulate the center/surround connectivity to enhance processing at the attended location and cause convergent RF shifts.

**Fig. S7.**
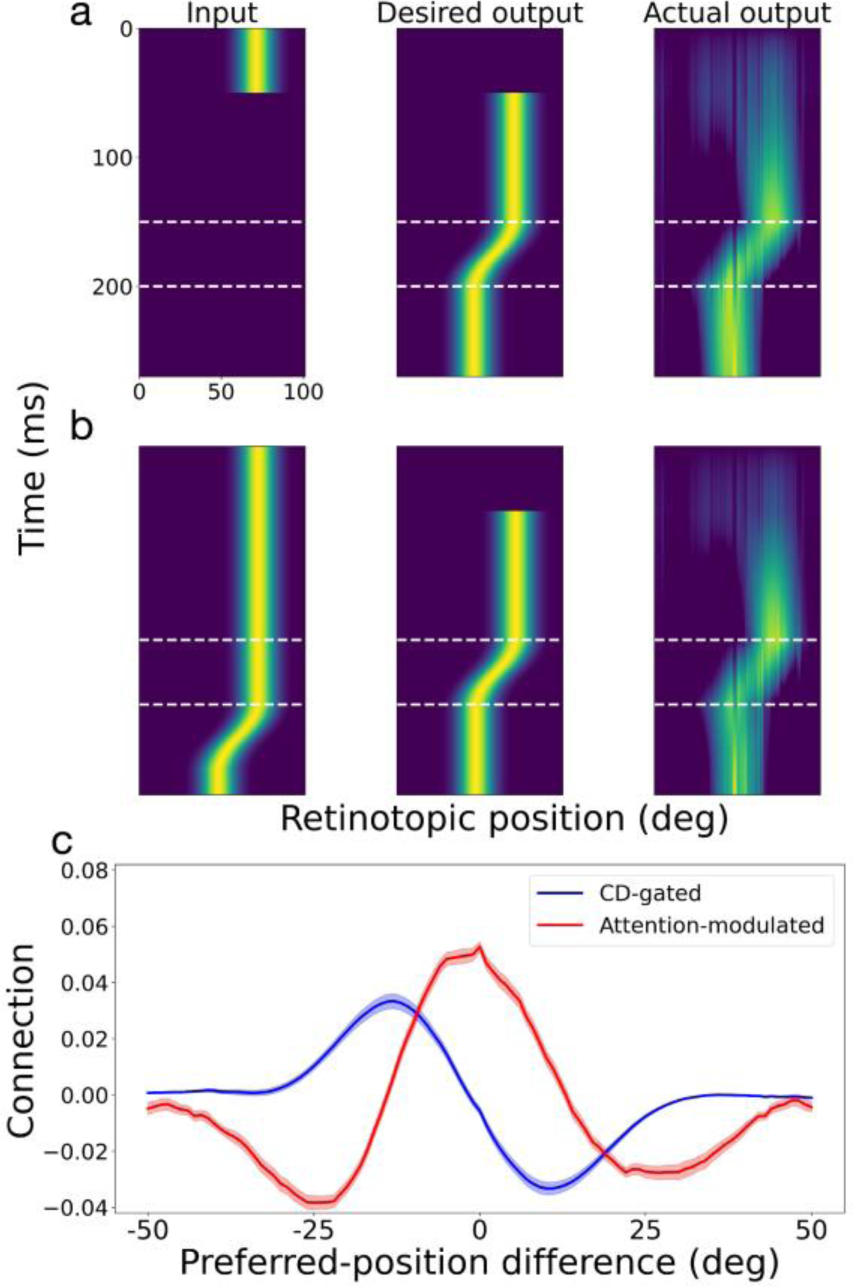
Automatic generation of the required connectivity patterns in the circuit model by training neural networks to predictively update retinal positions of both brief (a) and persistent (b) input stimuli during saccades. The format of the figure was identical to that for Fig. 10 of the main text except that both an example of brief input (a) and an example of the persistent input (b) are shown.

**Fig. S8.**
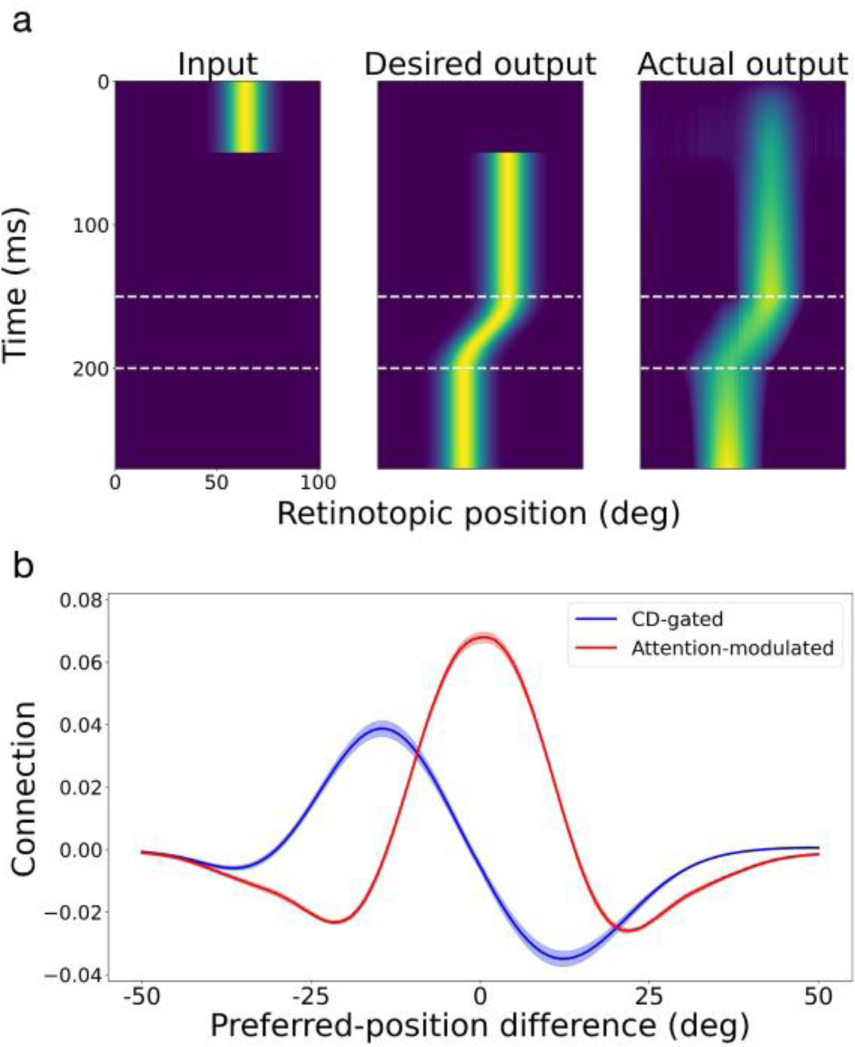
Automatic generation of the required connectivity patterns in the circuit model by training neural networks to predictively update retinal positions of brief input stimuli during saccades without attentional modulation. We still labeled the symmetric connections (red) as attention-modulated for easy comparison with Figs. 10 and S7. The format of the figure was identical to that for Fig. 10 of the main text.

## Acknowledgement

This work was supported by National Natural Science Foundation of China (32030045 and 32061143004), and US NIH (R01 EY032938) and NSF (1754211).

## Author contributions

MZ designed the experiments. NQ designed the models. MZ, NQ, and MEG supervised the project. CZ, LY, and MJ collected the data; XW, CZ, LY, and MJ analyzed the data. XW implemented and simulated the models. XW, LY, and MJ produced the figures. NQ, MEG, and MZ wrote the manuscript. All authors interpreted the data and edited the manuscript.

